# Evidence for ecological tuning of novel anuran biofluorescent signals

**DOI:** 10.1101/2023.07.25.550432

**Authors:** Courtney Whitcher, Santiago R. Ron, Fernando Ayala-Varela, Andrew Crawford, Valia Herrera-Alva, Ernesto Castillo-Urbina, Felipe Grazziotin, Randi M. Bowman, Alan R. Lemmon, Emily Moriarty Lemmon

**Affiliations:** Florida State University, Department of Biological Science, Tallahassee, 32306, USA; Museo de Zoología, Pontificia Universidad Católica del Ecuador, Escuela de Ciencias Bioloógicas, Quito, 170143, Ecuador; Universidad de los Andes, Department of Biological Sciences, Bogotá, 111711, Colombia; Museo de Historia Natural de la Universidad Nacional Mayor de San Marcos, Departamento de Herpetología, Lima, 15072, Perú; Instituto Butantan, Laboratório de Coleções Zoológicas, São Paulo, 05345, Brazil; Florida State University, Department of Scientific Computing, Tallahassee, 32306, USA

## Abstract

Our study assesses the variability of amphibian biofluorescence and provides insight into its potential functions and role in anuran evolution. Via a field survey across South America, we discovered and documented patterns of biofluorescence in tropical amphibians. We more than tripled the number of species that have been tested for this trait and added representatives from previously untested anuran families. We found evidence for ecological tuning (i.e., the specific adaptation of a signal to the environment in which it is received) of the novel anuran biofluorescent signals. Across groups, the fluorescence excitation peak matches the wavelengths most available at twilight, the light environment in which most frog species are active. Additionally, biofluorescence emission spans both wavelengths of low availability in twilight and the peak sensitivity of green-sensitive rods in the anuran eye, likely increasing contrast of this signal for a conspecific receiver. With evidence of tuning to the ecology and sensory systems of frogs, our results suggest frog biofluorescence is likely functioning in anuran communication.

## Introduction

Biofluorescence—an organism’s ability to absorb light and re-emit it at a longer wavelength (i.e., a biological organism’s ability to fluoresce; 1)—is present in a range of taxa including insects, plants, fishes, and reptiles (2) but was only recently discovered in amphibians (3). Across the tree of life, fluorescence has been found to signal sexual or resource attractiveness (bees and flowers (4), birds (5), spiders (6)), to contribute to species recognition (copepods (7)), to signal the condition of an organism (leaves and fruits (8–10), mammals (11)), and to facilitate camouflage (reef fishes (12)). Amphibian biofluorescence has been identified in multiple anuran families, but reports are taxonomically sparse, with descriptions of fluorescence in only a handful of species (13–19). Several studies have speculated on the function of fluorescence in amphibians, but empirical tests are lacking in the literature (16–19).

Biofluorescence has been proposed in amphibians to act as a visual signal to potential mates, predators, or other receivers (16–19). According to the sensory drive hypothesis, natural selection should favor signals that maximize the received signal relative to background noise (20–22). Thus, sensory drive should lead to evolutionary coupling of sensory systems, signals, signaling behavior, and habitat choice (20). With regard to visual signals such as biofluorescence, the coloration of an organism’s signal is predicted to depend on the spectrum of the ambient light in the environment (23). Unlike reflected coloration, fluorescent signals are dynamic because the chemicals that produce biofluorescence (fluorophores) manipulate the light present in the environment, absorbing light at one wavelength and re-emitting it at another wavelength (1). Hence, fluorescing organisms are not limited to only reflecting the color of light available in the environment. Variation in the excitation and emission colors of biofluorescence, with varying fluorophore chemical mechanisms, has evolved both across and within taxonomic groups (2, 24). The innate multidimensionality of biofluorescence may enhance evolutionary lability of this trait, enabling it to respond rapidly to selection within specific abiotic and biotic environments.

Since biofluorescence was discovered in frogs in 2017 (3), new accounts have documented the trait in all three orders of amphibians (18) and described emission patterns that range from body-wide (13–14) to location-restricted patches (15–18). These studies employed excitation light within the 365–460 nm range (ultraviolet, violet, or blue light), obtaining results that varied in the presence and intensity of any fluorescent signals, depending upon the light source, species, and particular study (13–18). Under this excitation range, species produced biofluorescent emissions in the 450–550 nm range (blue to green visual light (13–19)). Although studies have suggested that amphibian biofluorescence could increase perception of other conspecific individuals in low-light twilight environments (3,18), these predictions have not been tested.

Four criteria have been proposed for demonstrating that biofluorescence has a biological function, and therefore ecological significance (25). First, the fluorescent pigment will absorb the dominant wavelengths of light found in the environment. Second, the fluorescence will be viewed by the receiver against a contrasting background environment. Third, organisms viewing the fluorescence will have spectral sensitivity in the fluorescent emission range, allowing the fluorescence to be perceived. Finally, the fluorescent signals will be located on a part of the body displayed during signaling. Studies are lacking in any system, however, that directly test all four criteria (but see Lim et al. 2007 in jumping spiders (6) and Haddock and Dunn 2015 in siphonophores (26)), and a direct test for ecological significance of biofluorescence has yet to be completed for amphibians.

The goals of our study were to survey the phylogenetic breadth of anuran biofluorescence by increasing the number and taxonomic distribution of species sampled and to evaluate evidence for an association between environmental characteristics and biofluorescent emission across anurans. For the latter goal, we tested the four criteria proposed by Marshall and Johnsen (2017) for establishing the ecological significance of biofluorescence. By increasing the number of species in which the trait has been observed by more than 250%, our study provides deeper insight into the phylogenetic and functional significance of biofluorescence.

## Results

### Survey of biofluorescence

During the ten weeks of our field collections, we added biofluorescent measurements from representatives of one salamander family, one caecilian family, and 13 anuran families (Fig. 1). We increased the percentage of anuran families tested for biofluorescence from <17% to 24%, adding 39 genera and 151 species and increasing the percentage of genera and species tested from 5% to >8% and from 0.55% to 1.98%, respectively (Table 1). We more than tripled the number of species tested for this trait compared to those tested within the previous five years. We tested 528 individuals, quantifying biofluorescent emission in response to five different excitation light sources (previously only one or two of these light sources had been used in testing (13–18); Table 1). The new untested light sources frequently excited novel biofluorescent patterns that were missed in previous studies because the incorrect excitation wavelength was used. The previously tested species in which we revealed fluorescence by utilizing an additional excitation light source include *Boana cinerascens, Boana lanciformis, Dendropsophus rhodopeplus, Dendropsophus sarayacuensis, Phyllomedusa tarsius,* and *Phyllomedusa vaillantii* (16; Table 1).

**Figure 1.**
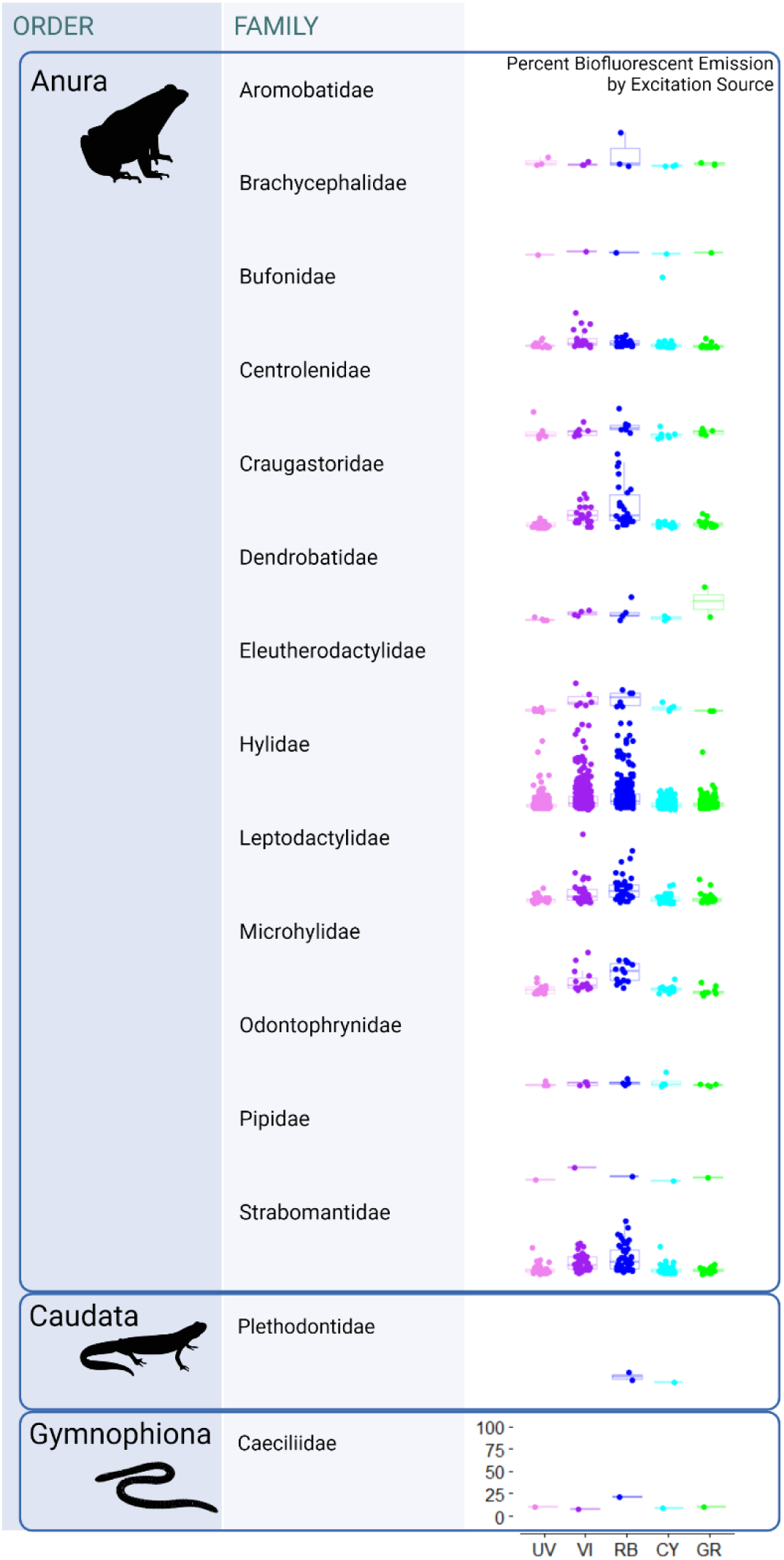
Biofluorescent emission by taxonomic family. A summary of the percent biofluorescent emission by family and excitation source. The set of box plots for each family presents the percent biofluorescent emission under the corresponding excitation light source: UV – Ultraviolet (360-380 nm), VI – Violet (400-415 nm), RB – Royal blue (440-460 nm), CY – Cyan (490-515 nm), and GR – Green (510-540 nm). Axis labels on the bottom panel hold for each set of box plots above. Each point on the plots represents one individual (the maximum percent biofluorescent emission recorded for that individual under that excitation light source). Each individual was measured under each light source. Percent biofluorescent emission refers to the percentage of total light shone on the individual that was re-emitted as biofluorescence. Table S1 contains the respective numeric values for these measurements.

**Table 1.**
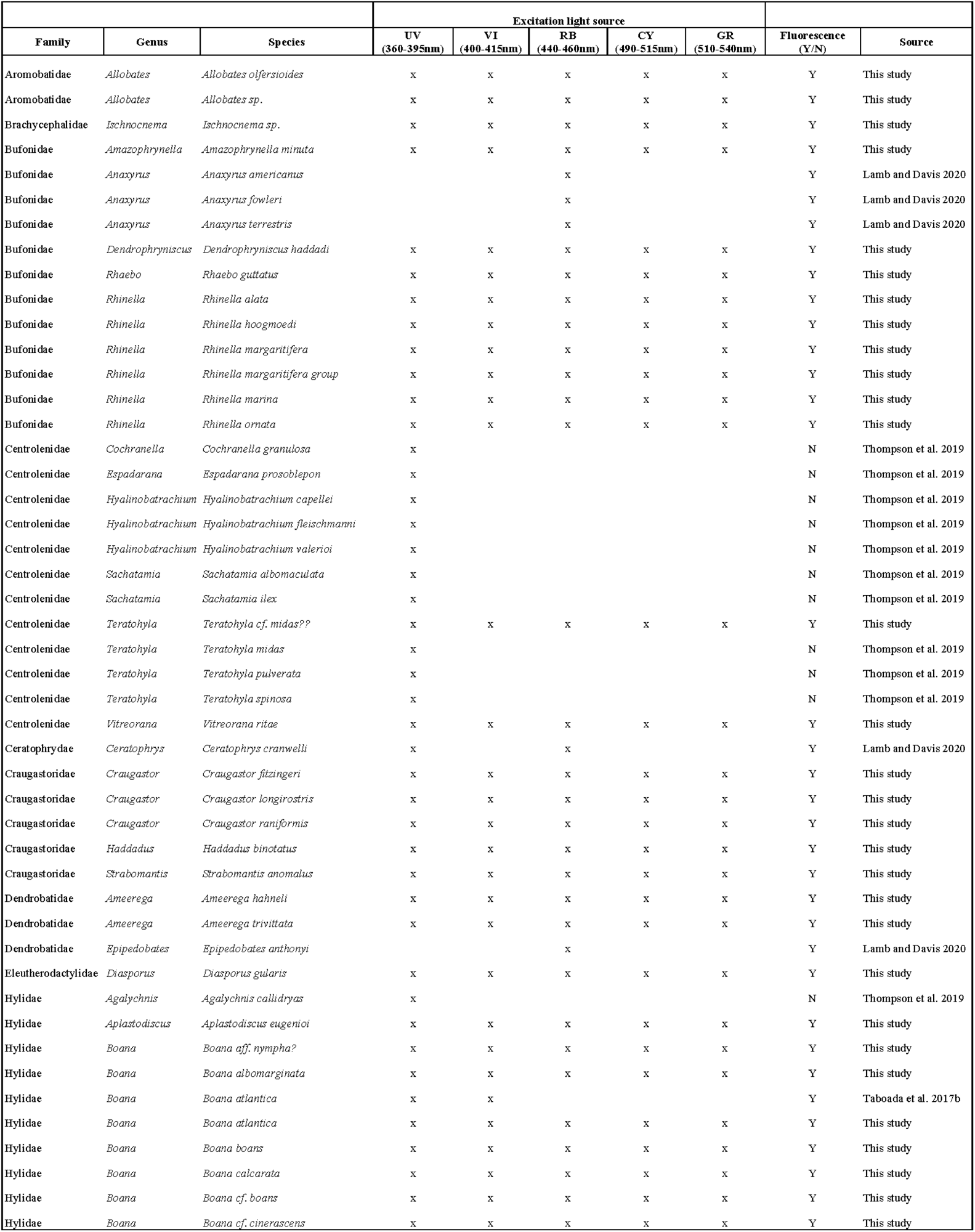
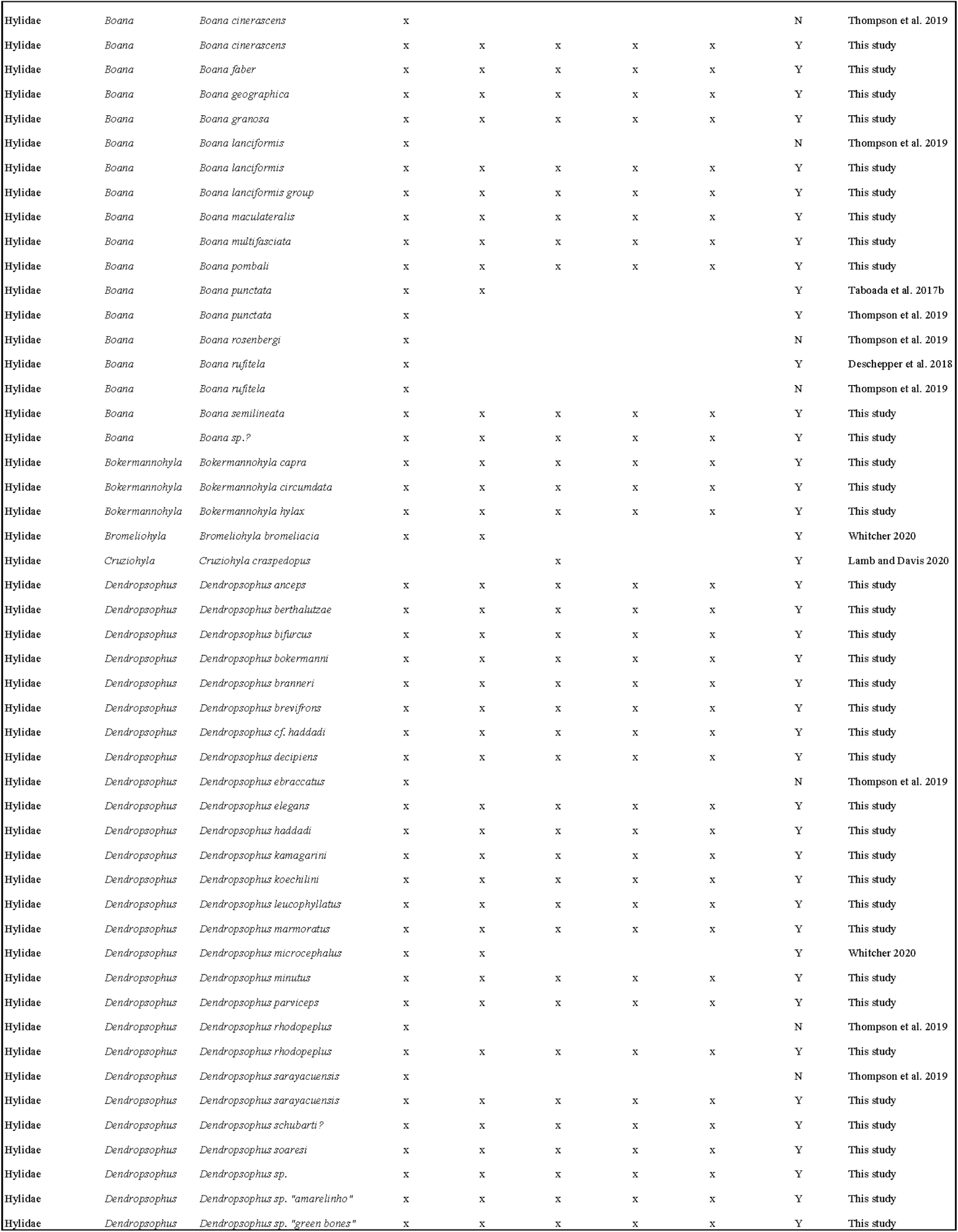
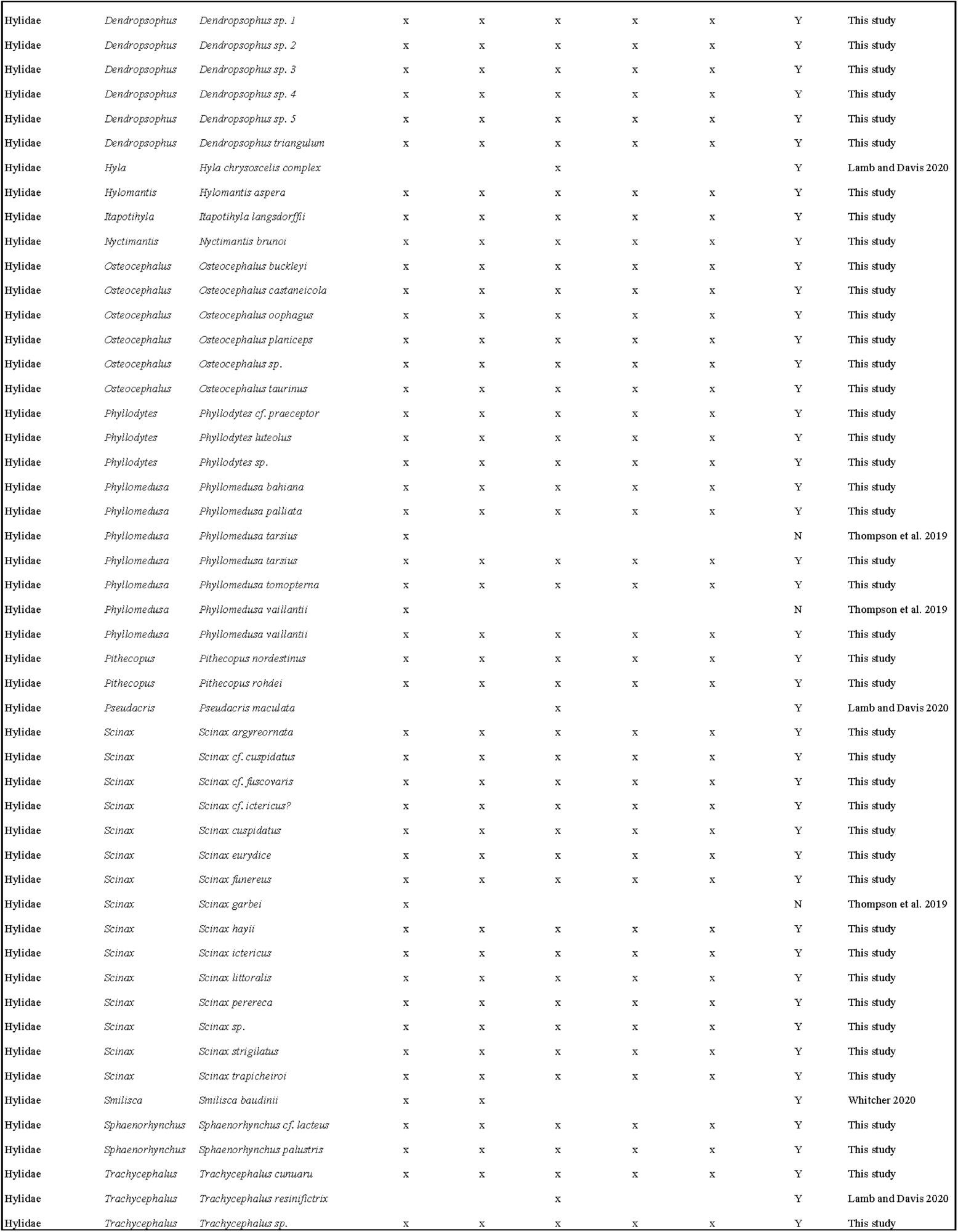
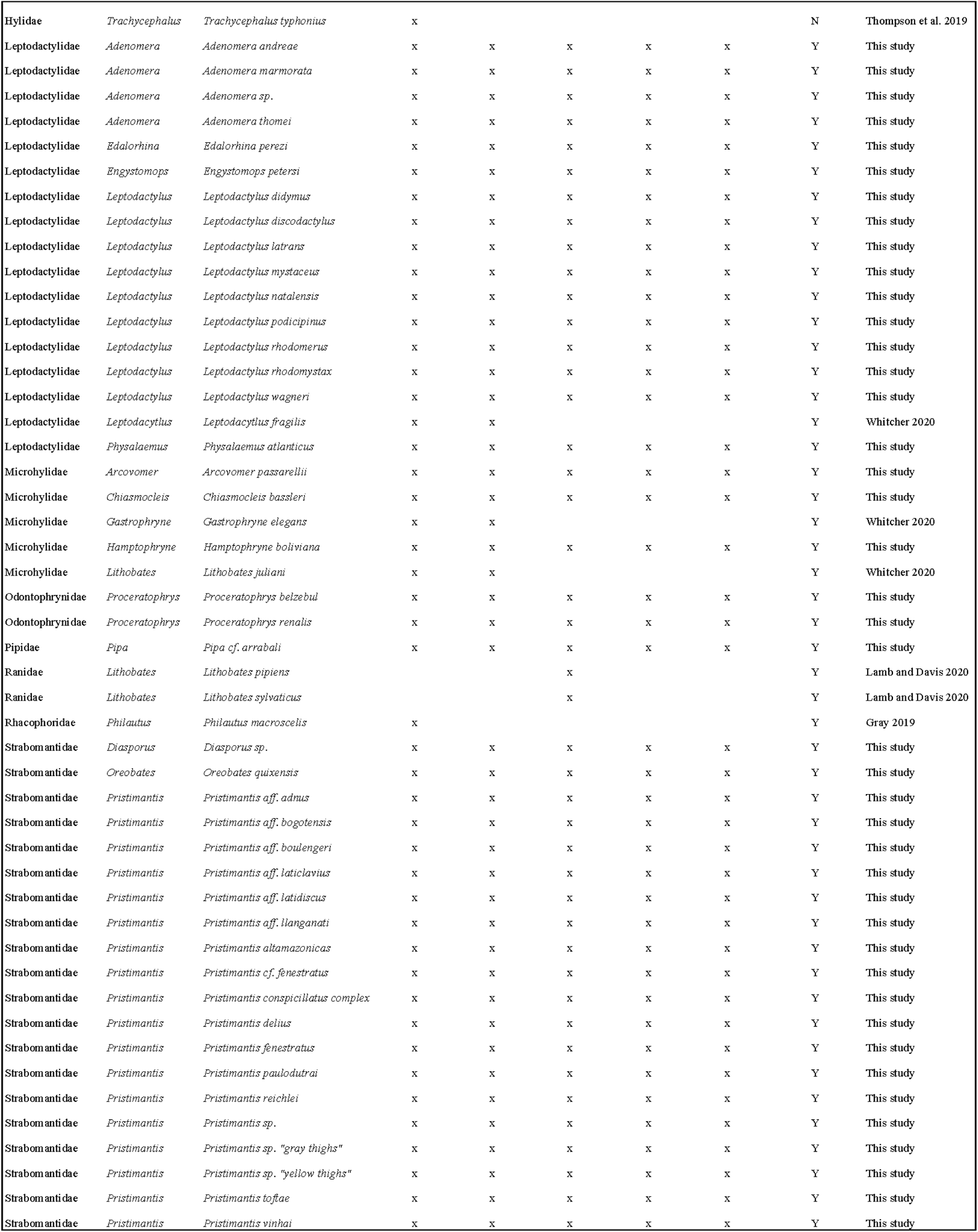
Results of previous and current studies testing anurans for biofluorescence. The excitation light source wavelength, resulting biofluorescence (Y/N), and source for documented tests of frog biofluorescence in the literature, by anuran family, genus, and species. Note the addition of excitation wavelengths in our study and our excitation of fluorescent signals in species already tested that were previously missed because the wrong excitation wavelength was used (see *Boana cinerascens, Boana lanciformis, Dendropsophus rhodopeplus, Dendropsophus sarayacuensis, Phyllomedusa tarsius,* and *Phyllomedusa vaillantii*).

### Assessing variation in biofluorescence

From each of our 17,692 spectrometer recordings we calculated a maximum percent biofluorescence emission (see Materials and Methods). The maximum percentage of biofluorescent emission from each individual under each excitation light source is presented by taxonomic family in Fig. 1 (Supp. Table 1). The maximum percentage of biofluorescent emission ranged from 1.95% to 96.85% with a mean of 11.11%.

We evaluated these maximum biofluorescence recordings against the predictions for each criterion proposed by Marshall and Johnsen (2017) for ecological significance (Fig. 2).

**Figure 2.**
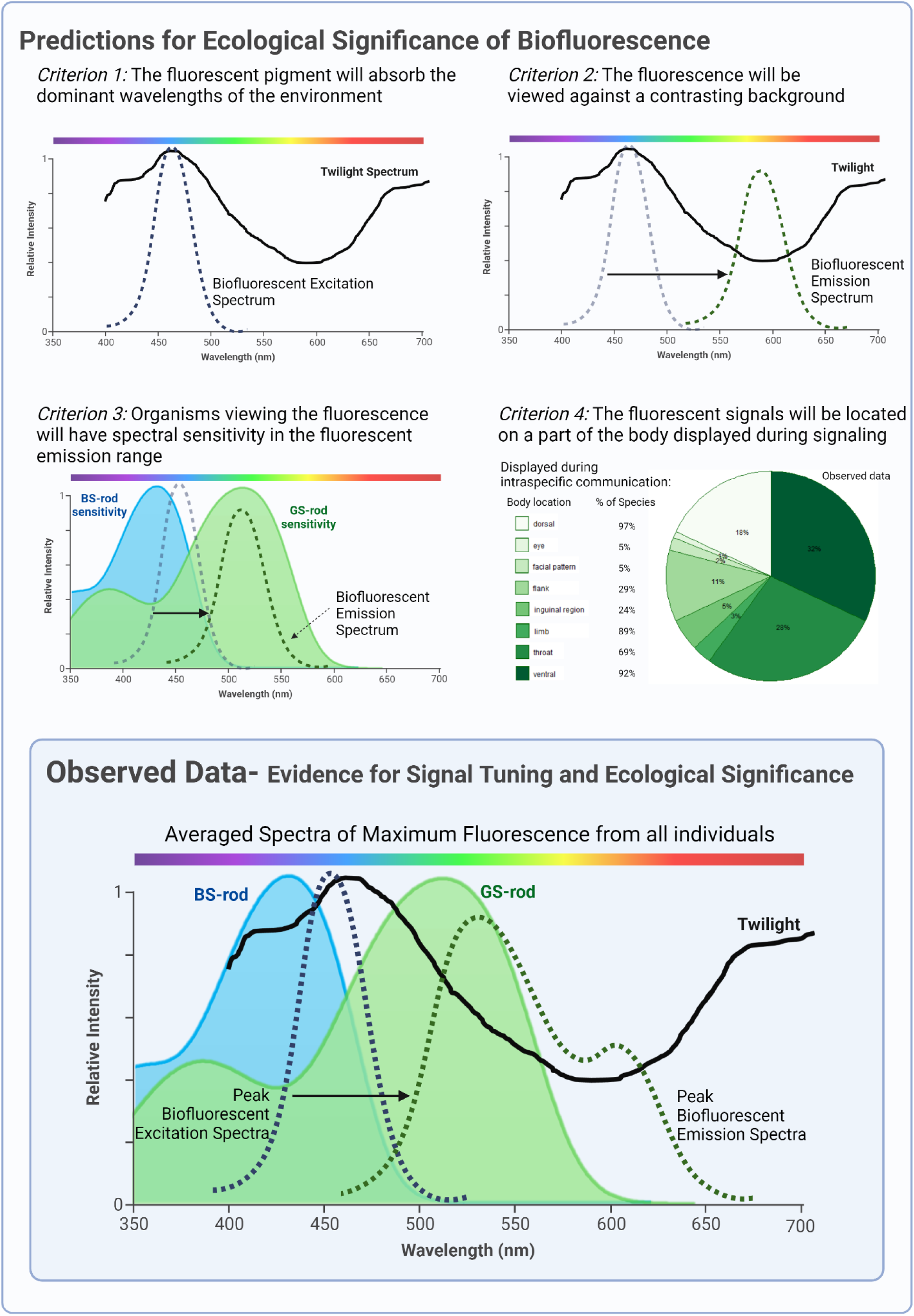
Predicted and observed evidence for ecological significance of biofluorescence. The top panel presents predicted results for each criterion proposed by Marshall and Johnsen (2017) that would suggest evidence of signal tuning and ecological significance of biofluorescence. The <u>predictions</u> under each criterion are presented within the framework of amphibian biology and ecology. *Criterion 1*: Peak fluorescence should be excited by absorbing the dominant wavelengths in the environment. In a twilight environment when most frogs are active (36), the dominant wavelengths are ∼450-460 nm (27; solid black line). The criterion predicts that peak excitation (blue dashed line) of anuran fluorophores should match this wavelength range. *Criterion 2*: Peak fluorescence emission (green dotted line) should be viewed against the least dominant wavelengths in the environment, to provide the most contrasting background. In a twilight environment, the least dominant wavelengths are ∼580-610 nm (27; solid black line). The arrow represents the Stokes Shift of the biofluorescence from peak absorption wavelengths to peak re-emission wavelengths. The criterion predicts the peak biofluorescence re-emission will be centered around ∼590 nm to provide the greatest contrasting background in a twilight environment. *Criterion 3*: Peak fluorescence emission (green dotted line) should match the spectral sensitivity of the intended receiver (green curve). When considering other anurans as the receiver, frogs have blue-sensitive (peak absorption ∼432 nm) and green-sensitive (peak absorption ∼500 nm) rods (28; 39). There are significantly more green-sensitive rods than blue-sensitive rods in the anuran retina (39). Hence, the criterion predicts the peak biofluorescence re-emission will be centered around ∼500 nm to match the greatest spectral sensitivity of another anuran receiver. *Criterion 4*: Peak fluorescence emission should be located on a part of the body displayed during signaling. Again, considering other anurans as the receiver, body locations displayed during *intraspecific* communication are listed (dorsal, eye, facial pattern, flank, inguinal region, limb, throat, and ventral). Of previously documented species that use intraspecific visual signals within the visual spectrum range, 97% have signal patterns on their dorsal surface, 92% on the ventral, 89% on the limb(s), 69% on the throat, 29% on the flanks, 24% on the inguinal region, 5% in the eyes, and 5% in the facial region (Fig. 2, Criterion 4; 29–34). The pie chart presents the observed data from our study of the body location from which the maximum biofluorescent recording from each individual was taken when the fluorescence was excited by blue (440-460 nm) light. The bottom panel presents the <u>observed data</u> for signal tuning and ecological significance from this study. When all fluorescent spectra recorded under blue excitation light (440-460 nm) are plotted (from all body locations), they follow the general shape presented by the dashed green line. This observed fluorescent emission pattern has peaks maximizing both the sensitivity of the green-sensitive rod in the anuran eye and the contrast with the background environment at twilight. The results from our study show that anuran biofluorescence meets all four criteria for ecological significance. Rod sensitivity spectra are from Yovanovich et al. 2017 (supp. mat.). The spectra of relative irradiance of wavelengths during twilight was digitized with permission from Cronin et al. 2014.

i. *Criterion 1:* The fluorescent pigment will absorb the dominant wavelengths of the environment.

We found support for this criterion: the excitation source with a peak wavelength closest to the dominant wavelength of the twilight environment produced the most fluorescence. The excitation wavelength that produces fluorescence is the wavelength of light absorbed by the fluorescent pigment; hence, the maximum fluorescent signal was produced by absorbing the wavelengths closest to those dominant in twilight. Twilight is defined as the light environment during the time between sunset and full night when the sun is between 0° and 18° below the horizon (27). We found a significant difference in biofluorescent emission intensity by excitation source (ꭓ2 = 446.88, p < 2.2e-16, n = 2,380). blue light excitation (440-460nm) produced a significantly greater percent fluorescent emission than any of the other excitation light sources (Table 2; Fig. 3). Additionally, the peak wavelength of the blue light excitation source (440-460 nm) is significantly closest to the dominant wavelength of the twilight environment (27; ꭓ2 = 2353.5, p < 2.2e-16, n = 2,353; pairwise comparisons Supp. Table 2). Although our sample size within each family is not large enough to statistically determine the influence of phylogeny on this result, the pattern is consistent across anuran groups (Fig. 1). The blue excitation light (440-460nm) induced fluorescence meets Criterion 1. Hence, we evaluate the subsequent criteria considering the fluorescent emission produced by this blue excitation light (440-460nm).

**Figure 3.**
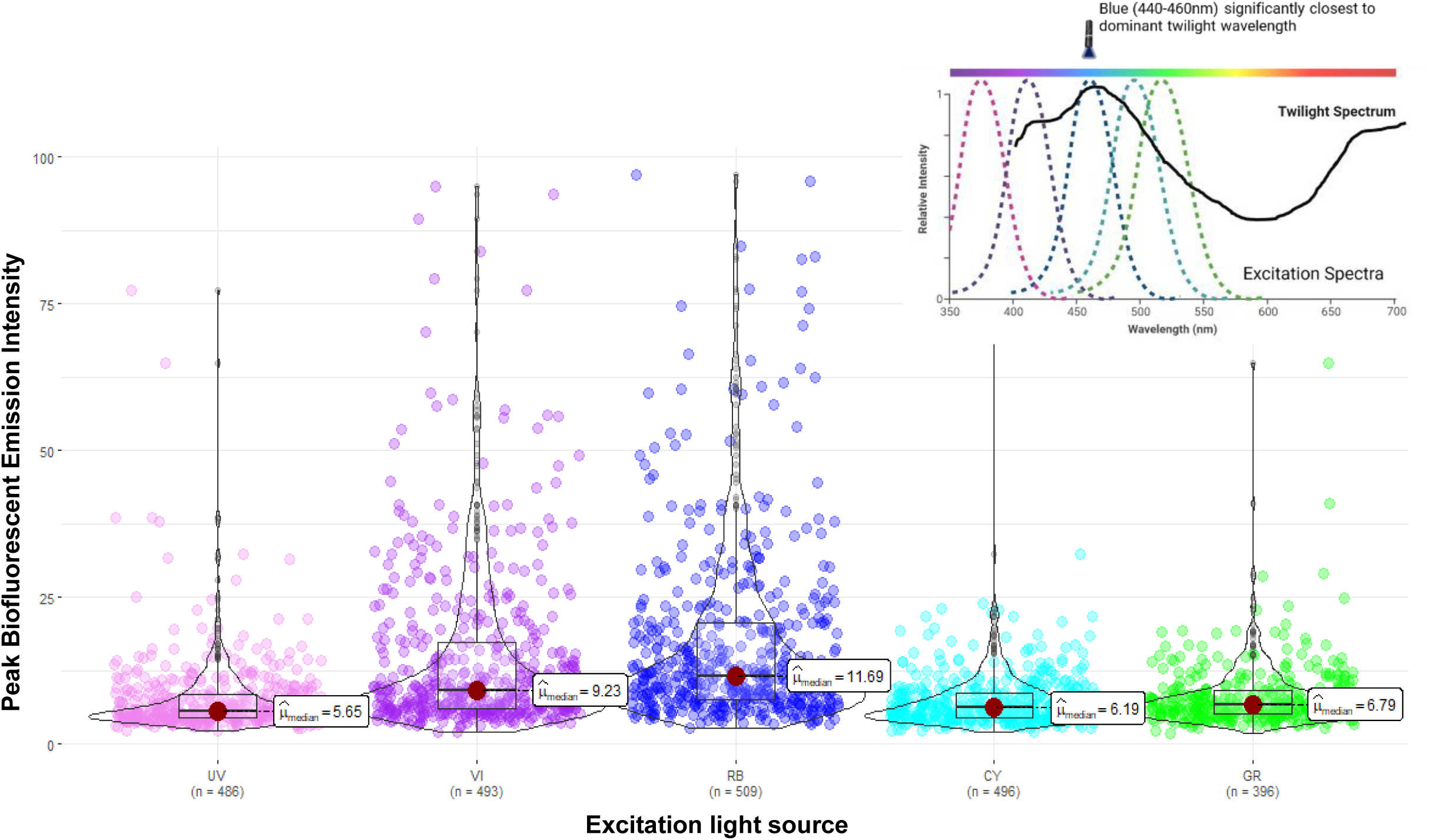
Biofluorescent emission intensity by excitation light source. The percent biofluorescent emission by excitation light source: UV – Ultraviolet (360-380 nm), VI – Violet (400-415 nm), RB – Royal blue (440-460 nm), CY – Cyan (490-515 nm), and GR – Green (510-540 nm). Each point represents one individual (the maximum percent biofluorescent emission recorded for that individual under that excitation light source). Percent biofluorescent emission refers to the percentage of total light shone on the individual that was re-emitted as biofluorescence. There is a significant difference in biofluorescent emission intensity by excitation source (ꭓ2 = 446.88, p = 2.05e-95, n = 2,380) with blue light excitation (RB, 440-460 nm) producing a significantly greater percent biofluorescent emission than any of the other excitation light sources. Additionally, the top right graph depicts that the blue (440-460 nm) light source has the closest excitation wavelength to the dominant wavelengths of the twilight environment (distance results presented in Supp. Table 2).

**Table 2.** Pair-wise comparison of percent biofluorescent emission by excitation light source. Dunn (1964) multiple comparison p-values adjusted with the Holm method for each excitation light source wavelength: UV – Ultraviolet (360-380 nm), VI – Violet (400-415 nm), RB – Royal blue (440-460 nm), CY – Cyan (490-515 nm), and GR – Green (510-540 nm). (*) Indicates significance at alpha = 0.05. All pairwise comparisons of percent biofluorescent emission are significantly different except for the biofluorescent emission produced by Ultraviolet and Green light. Blue (440-460 nm) excitation light produced the significantly highest fluorescence emission.

|  | UV | CY | GR | VI |
| --- | --- | --- | --- | --- |
| **CY** | 0.954<br>1.699e-01 |  |  |  |
| **GR** | 3.426<br>9.179e-04* | -2.538<br>1.115e-02* |  |  |
| **VI** | -12.274<br>4.346e-34* | 11.379<br>1.588e-29* | -8.190<br>6.536e-16* |  |
| **RB** | 16.965<br>7.409e-64* | -16.088<br>1.392e-57* | -12.596<br>8.885e-36* | 4.611<br>8.025e-06* |

ii. *Criterion 2:* The fluorescence will be viewed against a contrasting background.

Consistent with this prediction, the blue-light-induced fluorescent emission is viewed against a contrasting background in a twilight environment. The peak emission wavelengths overlap with the wavelengths of light least dominant in the twilight environment, providing the most contrast. We found a significant difference in biofluorescent emission wavelength by excitation source (ꭓ2 = 2021.46, p < 2.2e-16, n = 2,380; Supp. Fig. 1). Most notably, the peak emission wavelengths produced by blue light excitation (440-460nm) centered around approximately 527 nm and 608 nm (we refer to hereafter as a “green” peak and “orange” peak respectively; Fig. 4; Supp. Fig. 1). There were 192 individuals with a “green” emission peak and 316 individuals with an “orange” emission peak (Fig. 4). Both the “green” and “orange” fluorescent peak emission wavelengths match the least dominant wavelengths of the twilight environment better than expected by chance (p < 0.0001; Fig. 5). Hence, all fluorescent emission produced by blue light (440-460nm), which meets Criterion 1, also meets Criterion 2 by re-emitting fluorescence at wavelengths that provide the most contrast with the background environment.

**Figure 4.**
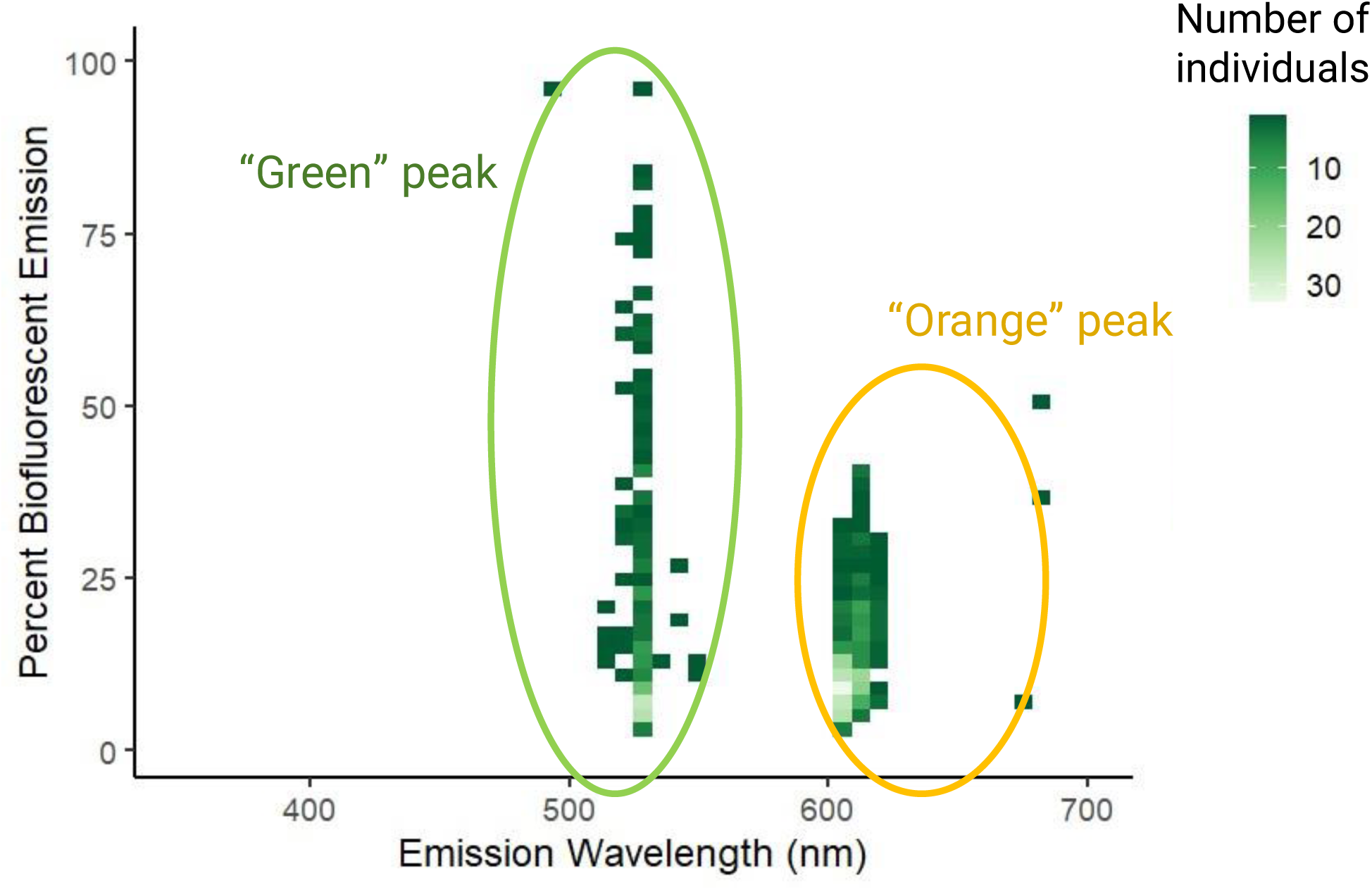
Peak biofluorescent emission wavelengths from blue (440-460 nm) excitation light. The percent biofluorescent emission by wavelength, where each point is colored by the number of individuals with a maximum percent biofluorescent emission recorded for that wavelength and intensity under the blue (440-460 nm) excitation light source. Lighter green corresponds to more instances of that wavelength and intensity emission. Note the peak emission wavelengths produced by blue light excitation are centered around ∼527 nm and ∼608 nm. Hence, we refer to these groupings of emission peaks as the “green” and “orange” peaks respectively.

**Figure 5.**
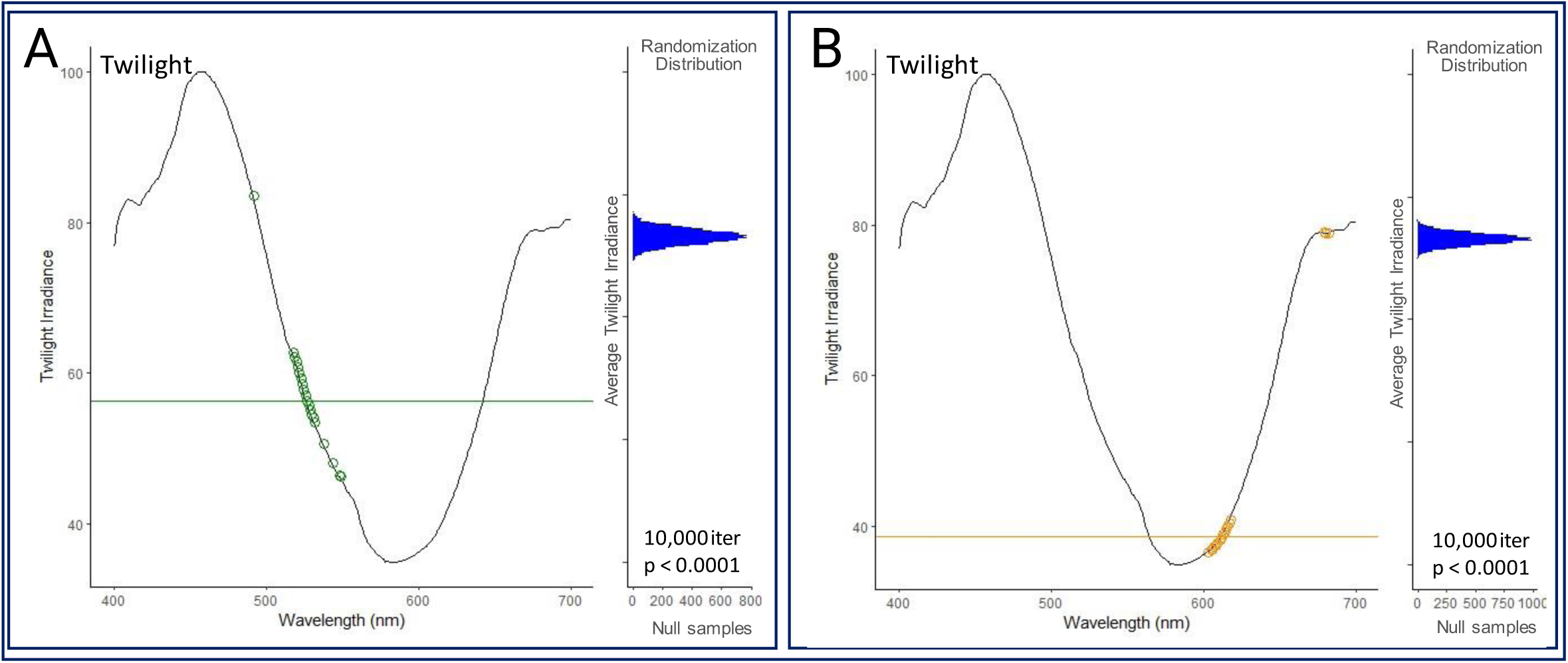
Randomization test results to evaluate Marshall and Johnsen (2017) Criterion 2: the fluorescence will be viewed against a contrasting background. Twilight irradiance spectrum (black line; digitized with permission from Cronin et al. 2014) compared to the “green” **(A)** and “orange” **(B)** fluorescence emission peaks. In each subplot, the observed wavelength of emission for different frogs are presented as colored circles. Total number of observed individuals is n = 192 and n = 316 for “green” and “orange” peaks respectively. To determine whether the average emission wavelengths (horizontal colored lines in each panel) are matched to the lowest twilight irradiance better than expected by chance, we compared those points to a null distribution. The null distribution was generated by sampling wavelengths uniformly between 400 and 700 nm and taking the average twilight irradiance (randomization distribution of test statistics presented as blue distribution on right in each panel). Each randomization distribution contains ten thousand iterations. P-values in the bottom right-hand corner of each graph present results of the comparison of the test statistic values in the randomization distribution to the observed test statistic. For both the “green” and the “orange” fluorescence emission peaks produced by blue excitation light, the peak fluorescent emission matched the wavelengths least available in the twilight environment better than expected by chance.

iii. *Criterion 3:* Organisms viewing the fluorescence will have spectral sensitivity in the fluorescent emission range.

Consistent with this criterion, the blue-light-induced “green” fluorescent emission closely matches the spectral sensitivity of the anuran green-sensitive rod. The emission wavelengths of the “green” peak overlap with the most sensitive wavelengths of light for the most abundant rod in the anuran visual system. As stated earlier, the peak emission wavelengths produced by blue light excitation (440-460nm) centered around approximately 527 nm and 608 nm, which we defined as a “green” and “orange” peak respectively (Fig. 4; Supp. Fig. 1). The “green” fluorescent peak emission wavelengths match the spectral sensitivity of the green-sensitive anuran rod better than expected by chance (p < 0.0001; Fig. 6); however, the “orange” fluorescent peak emission wavelengths do not (p > 0.9999; Fig. 6). Hence, the “green” peak fluorescent emission produced by blue light (440-460nm), which meets the first two criteria, also meets Criterion 3 by matching the spectral sensitivity of an anuran receiver in dim light.

**Figure 6.**
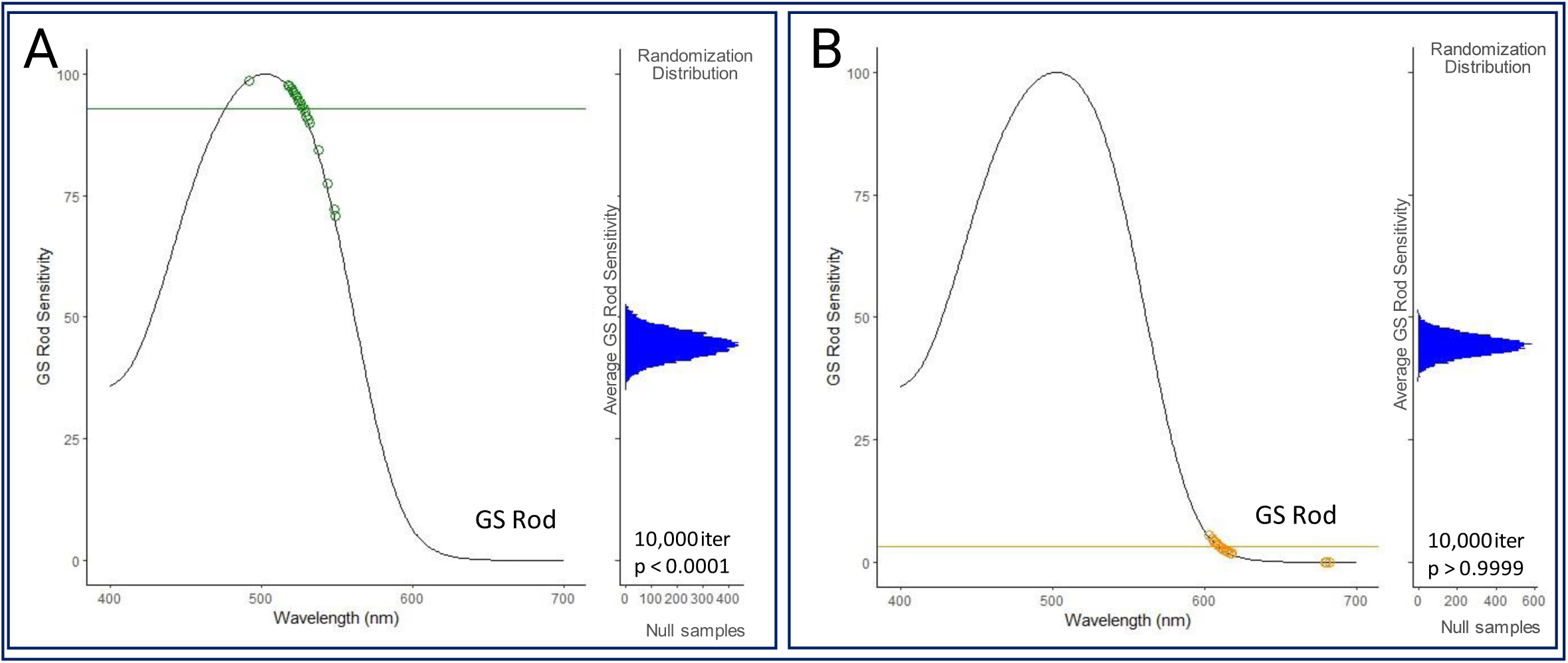
Randomization test results to evaluate Marshall and Johnsen (2017) Criterion 3: organisms viewing the fluorescence will have spectral sensitivity in the fluorescent emission range. Sensitivity curve of the green-sensitive (GS) rod of the anuran visual system (black line; obtained from Yovanovich et al. 2017) as compared to the “green” **(A)** and “orange” **(B)** fluorescence emission peaks. In each subplot, the observed wavelength of emission for different frogs are presented as colored circles on each plot. Total number of observed individuals is n = 192 and n = 316 for “green” and “orange” peaks respectively. To determine whether the average emission wavelengths (horizontal colored lines in each panel) are matched to the greatest sensitivity of the green-sensitive anuran rod better than expected by chance, we compared those points to a null distribution. The null distribution was generated by sampling wavelengths uniformly between 400 and 700 nm and taking the average GS rod sensitivity value (randomization distribution of test statistics presented as blue distribution on right in each panel). Each randomization distribution contains ten thousand iterations. P-values in the bottom right-hand corner of each graph present results of the comparison of the test statistic values in the randomization distribution to the observed test statistic. For the “green” fluorescence emission peak produced by blue excitation light, the peak fluorescent emission wavelengths matched the visual sensitivity of the green-sensitive anuran rod better than expected by chance. However, the “orange” fluorescence emission peak wavelengths did not.

Considering that both Criterion 2 and Criterion 3 must be met to interpret biofluorescence as an ecologically significant trait, we compared the blue-light-induced “green” emission peak to the tradeoff spectrum of twilight irradiance and anuran rod sensitivity. We divided the green-sensitive rod spectrum (28) by the twilight irradiance spectrum (27) to obtain the tradeoff spectrum of the wavelengths that would maximize both receiver sensitivity and background contrast (see Materials and Methods). The “green” fluorescent peak emission wavelengths match the tradeoff spectrum better than expected by chance (p < 0.0001; Fig. 7). Hence, these fluorescent emissions maximize the conditions of Criteria 2 and 3 simultaneously.

**Figure 7.**
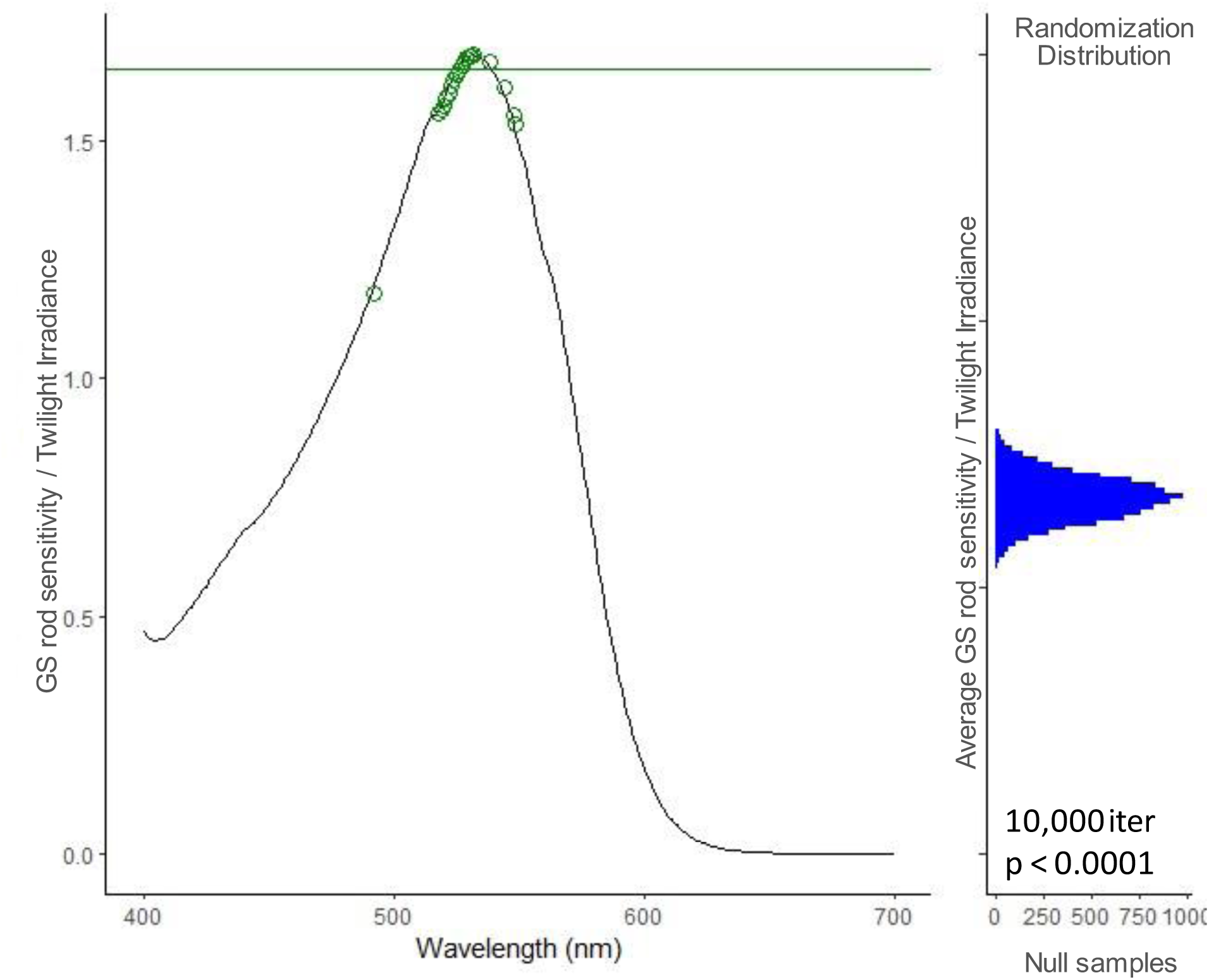
Randomization test results to evaluate meeting both Criterion 2 and Criterion 3 proposed by Marshall and Johnsen (2017). The black line represents the tradeoff between visual sensitivity and background contrast (green-sensitive rod sensitivity curve divided by twilight irradiance curve). The observed average tradeoff value (colored line) and the randomization distribution of test statistics (blue distribution) for “green” emission peak produced by blue (440-460 nm) excitation light. Total number of samples: n = 192 (colored points on graph). The randomization distribution contains ten thousand iterations. The p-value in the bottom right-hand corner presents results of the comparison of the test statistic values in the randomization distribution to the observed test statistic. The “green” fluorescence emission peak wavelengths produced by blue excitation light match the tradeoff between the visual sensitivity of the green-sensitive anuran rod and the contrast with the twilight environment better than expected by chance.

iv. *Criterion 4:* The fluorescent signals will be located on a part of the body displayed during signaling.

Consistent with this prediction, we found biofluorescence on regions of the body often displayed during intraspecific signaling, such as the dorsal surface and vocal sac, but location varied across species (see Discussion). The maximum biofluorescent emission for most individuals was produced under blue (440-460 nm) excitation light (261 of 512 individuals; Fig. 8). Within this subset of individuals, 32% of the maximum biofluorescent emission recordings came from a ventral pattern, 28% from the throat, 18% from a dorsal pattern, 11% from the flank, 5% from the inguinal region, 3% from a limb, 2% from a facial pattern, and 1% from the eye (Fig. 2; Supp. Table 3). For this criterion, we considered the maximum fluorescent emission recording from each individual under any excitation light source, as this insured that only one body location per individual was considered (see Materials and Methods). Though examining the maximum biofluorescent emission recording for each individual under each excitation light source (as in the analyses above) produced similar resulting percentages of fluorescence by body location (Supp. Table 4). Within these body regions, there was great variation in the pattern of biofluorescence (Fig. 8). Dorsal biofluorescence could result from secretions from the frog skin (as in *Boana atlantica*) or be present in distinct locations (as in *Hamptophryne boliviana* and *Scinax strigilatus*). Ventral biofluorescence could be widespread, condensed to specific patterns, or scattered in a speckled pattern (as seen in *Boana geographica, B. lanciformis,* and *Proceratophrys renalis*, respectively). Additionally, ventral biofluorescence often showed both green and orange emission (∼527 nm and ∼608 nm; as seen in *B. geographica* and *P. renalis*). Finally, biofluorescence could be present in distinct regions of the frog body, such as the forelimbs, throat, or eyes (as seen in *Chiasmocleis bassleri, Scinax trapicheiroi,* and *B. calcarata* respectively). Many of the body locations in which fluorescence was found are displayed during intraspecific signaling (Fig. 2). Of previously documented species that use intraspecific visual signals within the visual spectrum range, 97% have signal patterns on their dorsal surface, 92% on the ventral, 89% on the limb(s), 69% on the throat, 29% on the flanks, 24% on the inguinal region, 5% in the eyes, and 5% in the facial region (Fig. 2, Criterion 4; 29–34). Three of these locations are also the most common biofluorescent areas found in species we tested (dorsal, ventral, and throat; Fig. 2; Supp. Table 3), a finding that is consistent with Criterion 4.

**Figure 8.**
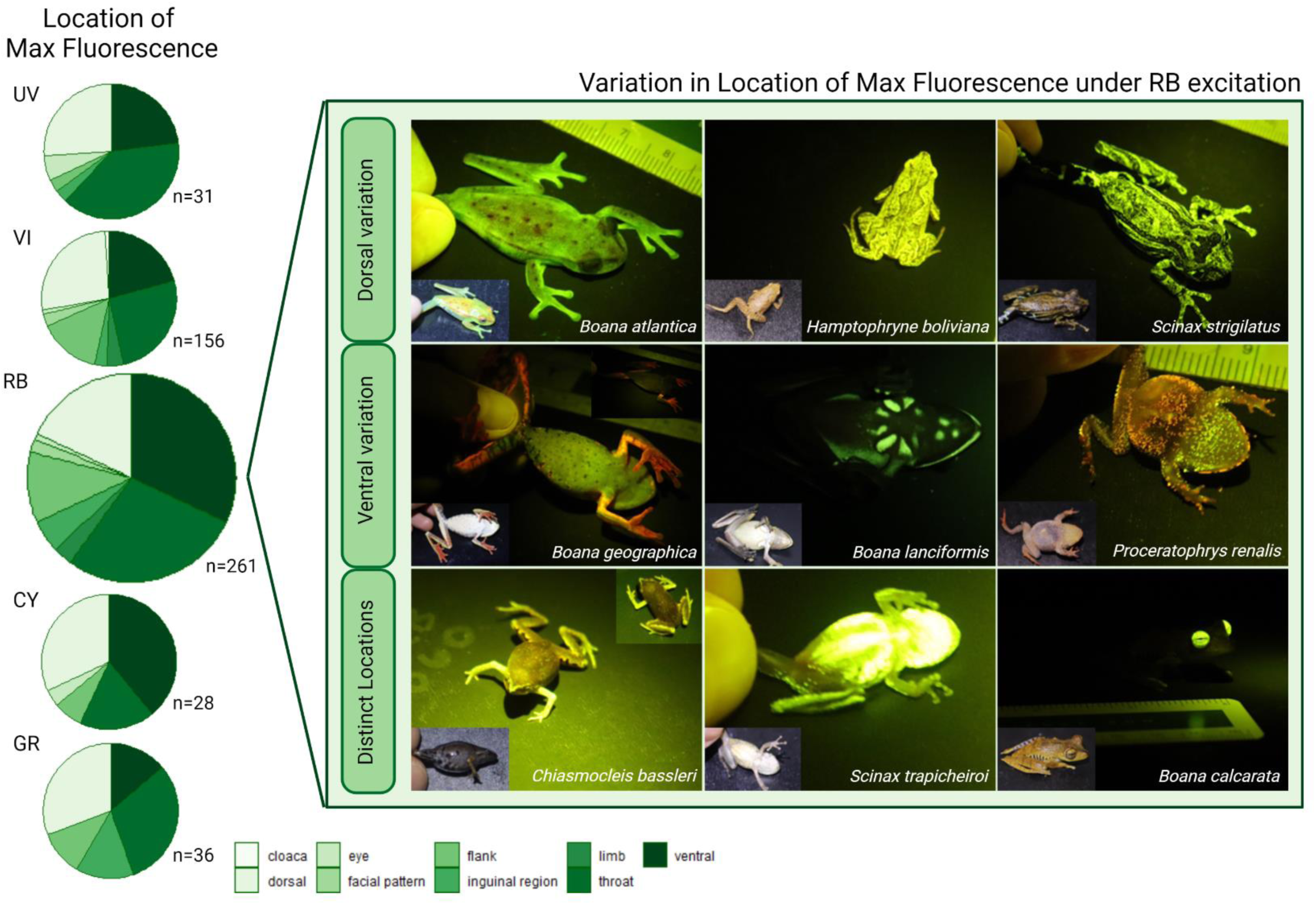
Maximum biofluorescence by body location. The left panel presents the body location from which the maximum biofluorescent emission recording from each individual was taken. Body regions were summarized into the following nine groups: cloaca, dorsal (including spectrometer recordings with a body location specified from any dorsal pattern), eye, facial pattern (including lip, spots under the eye, snout, etc.), flank, inguinal region, limb (including forelimb, thigh, etc.), throat (including vocal sac), and ventral (including any ventral pattern). The right panel shows some of the variation seen in the patterns of biofluorescence produced by blue (440-460 nm) excitation light. The species photographed, in order from left to right are: (top) *Boana atlantica, Hamptophryne boliviana, Scinax strigilatus*, (middle) *Boana geographica, Boana lanciformis, Proceratophrys renalis,* (bottom) *Chiasmocleis bassleri, Scinax trapicheiroi, and Boana calcarata*. Each photo panel has a photograph of the individual taken under blue (440-460 nm) excitation light through a 500 nm longpass filter and a photograph of the same individual taken under a full spectrum light source (inset). Dorsal biofluorescence could be produced by secretions from the frog skin (as in *Boana atlantica*) or present in distinct locations (as in *Hamptophryne boliviana* and *Scinax strigilatus*). Ventral biofluorescence could be widespread, condensed to specific patterns, or scattered in a speckled pattern (as seen in each individual of the middle row respectively). Additionally, ventral biofluorescence often showed both green and orange emission (∼527 nm and ∼608 nm; as seen in *Boana geographica* and *Proceratophrys renalis*). Finally, distinct regions of the frog body, such as the arms, throat, or eyes sometimes produced the greatest biofluorescent emission recording from an individual (as seen in each individual of the bottom row respectively).

## Discussion

### Survey of biofluorescence

Over a ten-week period, we more than tripled the number of frog species tested for fluorescence within the previous five years. Our sampling and results highlight the importance of testing additional amphibian species for biofluorescence and doing so under a wider range of excitation light sources. We added biofluorescence data for 528 individuals (representing 153 amphibian species) under five excitation light sources, when previously at most two had been used (13–18; Table 1). With the addition of these extra excitation wavelengths, we often excited fluorescent signals in species already tested that were previously missed because the incorrect excitation wavelength was used. The previously tested species in which we revealed fluorescence by utilizing an additional excitation light source include *Boana cinerascens, Boana lanciformis, Dendropsophus rhodopeplus, Dendropsophus sarayacuensis, Phyllomedusa tarsius,* and *Phyllomedusa vaillantii* (16; Table 1). The additional biofluorescence we revealed in our study highlighted important patterns in the evolution of this trait.

### Assessing variation in biofluorescence

*Criterion 1:* The fluorescent pigment will absorb the dominant wavelengths of the environment.

The excitation wavelengths that produce the most fluorescence are those wavelengths most abundant at the time of day when frogs are active. We found a significant difference in biofluorescent emission intensity by excitation source, with blue light excitation (440-460nm) producing significantly greater percent fluorescent emission than any of the other excitation light sources (Table 2; Fig. 3). Additionally, the peak excitation of the blue light excitation source is significantly closest to the dominant wavelength of the twilight environment (27; Supp. Table 2). In a twilight environment, wavelengths of blue light are at a higher relative abundance than any other wavelengths in the environment (27, 35; twilight spectra in Fig. 5 digitized with permission from Cronin et al. 2014). Additionally, most frogs are nocturnal and active during these twilight hours (36). Evidence shows that the light environment plays an important role in shaping animal coloration and the evolution of signals (37, reviewed in 22). Hence, the match between the peak excitation wavelength (the wavelengths that produce the most fluorescence) and the wavelengths most available in the environment at the time of day when frogs are active is notable. The sensory drive model predicts that environmental constraints will drive the evolution of a signal to match both the environmental transmission properties in the habitat and the sensory biases of the receiver in that habitat (22). Here, we found evidence that the excitation of anuran fluorescence matches the dominant wavelengths available in the environment.

The blue (440-460nm) excitation light induced fluorescence meets Criterion 1. Hence, we evaluate the subsequent criteria considering the fluorescent emission produced by this blue (440-460nm) excitation light.

*ii. Criterion 2:* The fluorescence will be viewed against a contrasting background.

We found that the blue-light-induced fluorescent emission is viewed against a contrasting background in the twilight environment. The peak emission wavelengths match the wavelengths of light least dominant in the twilight environment, providing the most contrast. We found a significant difference in biofluorescent emission wavelength by excitation source (Supp. Fig. 1). Most notably, the peak emission wavelengths produced by blue light excitation (440-460nm) centered around 527 nm and 608 nm (Fig. 4; Supp. Fig 1.). In a twilight environment, wavelengths of blue light are at the highest relative abundance and wavelengths of orange light (∼590-620nm) are at the lowest relative abundance compared to any other wavelengths in the environment (27, 35; twilight spectra in Fig. 5 digitized with permission from Cronin et al. 2014). Signals should be most easily detected when they differ from the background environment (38). The observed re-emission of both the “green” and “orange” fluorescent peak wavelengths match the least dominant wavelengths of the twilight environment better than random (Fig. 5). Anuran biofluorescence is absorbing light at the wavelengths most abundant in the environment and re-emitting it at the wavelengths least abundant in the environment. Hence, biofluorescence is increasing the contrast of the frog visual signal in a twilight environment, as compared to the reflected color of an individual without fluorescent properties in the same environment.

*iii. Criterion 3:* Organisms viewing the fluorescence will have spectral sensitivity in the fluorescent emission range.

Peak biofluorescent emission matches the peak sensitivity of the anuran eye. We found that the blue-light-induced fluorescent emission matches the spectral sensitivity of the anuran green-sensitive rod. The emission wavelengths of the “green” peak closely match the most sensitive wavelengths of light for the most abundant rod in the anuran visual system.

Cones help organisms see in bright light and rods help organisms see in dim light (39). Frogs have two rods that allow for color vision in dim light, and most anuran species mate in dim light, twilight conditions. Frogs have blue-sensitive (peak absorption ∼432nm) and green-sensitive (peak absorption ∼500nm) rods (28, 39; sensitivity curves of blue-sensitive and green-sensitive rods in Fig. 4 obtained from Yovanovich et al. 2017). There are significantly more green-sensitive rods than blue-sensitive rods in the retina of frogs (39). Specifically, Denton and Wyllie (1955) found that green-sensitive rods occupy about 60% of the retinal area while blue-sensitive rods occupy only about 8% of the retina in *Rana temporaria* (39).

We found evidence that the individuals with a “green” fluorescent peak emission wavelength match the spectral sensitivity of the green-sensitive anuran rod better than random (Fig. 6). The re-emission of light to a longer wavelength via fluorescence pushes the amphibian color emission to the peak sensitivity of the most abundant green-sensitive rods. Yovanovich and colleagues (2017) found specific amphibian phototaxis preference of green over blue signals only under the dim lighting conditions in which frogs are active. Hence, re-emitting blue light as green, under dim-light conditions, increases the visual signal of the frog to other frog receivers. Violet-induced fluorescence contributed nearly 30% to the total emerging light under twilight conditions in *Boana punctata* (3). As violet light produced the second highest biofluorescent emission in our study, only behind blue light, the contribution of blue-light-induced anuran biofluorescence in twilight conditions is likely even higher. Our evidence of a match between the anuran fluorescent signal and anuran optical sensitivity suggests biofluorescence is increasing the visual signal of frogs to conspecifics. The evidence of ecological and spectral tuning of anuran biofluorescence suggests this trait likely functions in frog communication and/or behavior.

The individuals with an “orange” fluorescent peak emission wavelength, however, do not match the spectral sensitivity of the green-sensitive anuran rod better than random (Fig. 6).

This finding could suggest that orange fluorescence may be a byproduct of the evolution of a fluorophore with a green and orange peak, as there were multiple emission spectra in our dataset with both peaks. Additional analysis of the specific fluorophore chemical producing the biofluorescence across species would be needed to test this hypothesis. Additionally, the orange fluorescent emission could serve as a signal to a different intended receiver (e.g., a predator or other heterospecific viewer of the signal). We evaluated the criteria for ecological significance of biofluorescence within the context of the biology and ecology of other anuran receivers; however, the visual sensitivities of *hetero*specific receivers (e.g., a predator or prey viewer of the signal) and the environments in which these types of interspecific communications occur vary greatly (27). Further examination of whether our data meet these criteria within the context of different environments and different intended receivers would aid in evaluating other potential mechanisms of the fluorescent signals recorded.

The “green” peak fluorescent emission produced by blue light (440-460nm) meets the criteria for ecological significance of biofluorescence with respect to a conspecific receiver. Specifically, considering that both Criterion 2 and Criterion 3 must be met to support the ecological significance of this trait, we compared the blue-light-induced “green” emission peak to the tradeoff spectrum of twilight irradiance and anuran rod sensitivity. The individuals with a “green” fluorescent peak emission wavelength match the tradeoff spectrum better than random (Fig. 7). Hence, the fluorescent emission of these individuals maximizes Criterion 2 and 3 simultaneously. In 192 individuals spanning 87 species, anuran biofluorescence absorbs the wavelengths most dominant in the twilight environment and re-emits that light to match the tradeoff between visual sensitivity and background contrast, maximizing both. Additionally, this green emission peak produces the most intense fluorescence, re-emitting up to nearly 97% of the light shone on the frog in some cases (Fig. 4). This observed evidence of non-random excitation and emission wavelengths suggests the green fluorescence induced by blue light is likely functioning in anuran communication in these species.

*iv. Criterion 4:* The fluorescent signals will be located on a part of the body displayed during signaling.

In addition to exploring the patterns of ecological and spectral tuning of anuran biofluorescence across taxonomic families, we also examined variation in this signal upon which selection could act. The maximum percentage of biofluorescent emission ranged from 1.95% to 96.85%, and the body location that produced the greatest biofluorescent emission for each individual varied. Figure 8 shows some of the diversity in fluorescent pattern observed. Dorsal biofluorescence could be produced by secretions from the frog skin (as in *Boana atlantica*) or present in distinct locations (as in *Hamptophryne boliviana* and *Scinax strigilatus*). Ventral biofluorescence could be widespread, condensed to specific locations, or scattered in a speckled pattern (as seen in *Boana geographica, B. lanciformis,* and *Proceratophrys renalis* respectively). Additionally, ventral biofluorescence often showed both green and orange emission (∼527nm and ∼608nm; as seen in *B. geographica* and *P. renalis*). Finally, distinct regions of the frog body, such as the forelimbs, throat, or eyes sometimes produced the greatest biofluorescent recording from an individual (as seen in *Chiasmocleis bassleri, Scinax trapicheiroi,* and *Boana calcarata* respectively).

Many of the parts of the body in which fluorescence was found are displayed during intraspecific signaling in nature (Fig. 2). The percentage of documented visually-communicating anuran species grouped by body region of the visual signal is shown in Fig. 2, Criterion 4. Overall, 97% of species display the dorsal, 92% ventral, 89% limb(s), 69% throat, 29% flank, 24% inguinal region, 5% eye, and 5% facial pattern (of 62 species; 29–34). Additionally, numerous species display multiple regions of the body, potentially sending different information to different receivers (e.g., to attract females or to deter rival males; 29–34). In this study, many of the bodily locations often displayed during intraspecific signaling in nature produced a fluorescent signal. For example, the male *Dendropsophus parviceps* individual we tested had its maximum fluorescence located on the lateral flank region and induced by blue light (96.85% emission intensity). This species is known to have multiple intraspecific visual displays including toe trembling, arm waving, foot flagging, and throat displays (33). This fluorescent flank region of the body is specifically presented during the limb waving displays; hence, the green fluorescent emission of this body region is likely contributing to the visual signal of these intraspecific displays in dim light. Increased sampling within each species and further species-specific examination is needed to evaluate the contribution of fluorescence to intraspecific visual displays, since the body locations utilized in intraspecific signaling vary widely across and within anuran groups.

The fluorescent locations and patterns we observed in our study can provide insight into potential functions of biofluorescence in anurans. For example, dorsal or facial fluorescent patterns could be employed for species recognition in certain groups (such as in stomatopods (40) and proposed in reef fishes (41)). In addition, fluorescent throat surfaces, which represent the brightest body region in 28% of our study individuals, could be used for mate choice or species recognition via male vocal sac movement during calling (blue-light-induced fluorescence; Fig. 8; Supp. Table 2). The role of fluorescent signals in mate choice in other taxa (specifically in budgerigar parrots (5) and jumping spiders (6)) and recent findings of sexual dimorphism in amphibian fluorescence (19) make this a likely function worthy of exploration. Fluorescence in the inguinal (inner thigh) region, normally invisible when the frog is at rest, could serve to startle a potential predator while its prey escapes (42). Additionally, the relatively low occurrence of biofluorescence in the inguinal region could be attributed to the possibility that these color patterns are primarily associated with anti-predator coloration serving as a form of *interspecific* rather than *intraspecific* communication.

In some taxa, patterns of biofluorescence appear to be evolutionarily conserved. For example, species from two genera of microhylids have similarly fluorescent arms: *Chiasmocleis bassleri* (this study; Fig. 8) and *Gastrophryne elegans* (17). In other taxa, biofluorescence trait evolution is much more labile, as seen in the extreme pattern and location variation within the hylid genus *Boana*. Specifically, we found that *B. atlantica* has intensely fluorescent secretions, *B. geographica* has an elaborate fluorescent dorsal pattern, *B. lanciformis* has multi-colored emission of the ventral surfaces and limbs, and *B. calcarata* has intense fluorescence of the eyes/irises (Fig. 8). The one comparative study on the evolution of biofluorescence examined coarse scale evolution of the trait across genera of varying taxonomic groups (43). As the many missing taxa are characterized with further work, a clearer picture of the evolutionary lability of biofluorescence will emerge.

### Caveats

Further examination of the biofluorescent signal in anuran groups spanning different ecological niches is needed to assess if these patterns hold. While our sample sizes within Dendrobatidae and Pipidae are extremely small and not statistically robust enough to test, the pattern of peak biofluorescence being excited by blue light (440-460nm) may not hold in these two families (Fig. 1). As these are largely diurnal and aquatic families respectively, the environmental spectra, transmission, and hence the resulting wavelengths available for producing a fluorescent signal differ drastically from the rest of the Anura families (27, 35). Additionally, the green-sensitive rod spectra utilized in our analyses was obtained exclusively from two families, Ranidae and Bufonidae, which are known to be primarily nocturnal. Future work should compare the maximum biofluorescent emission wavelength to species specific spectral sensitivities as that data becomes available. Recent evidence suggests that ecology and diurnality shape the visual sensitivities of frog vision (44–45). Hence, examining how biofluorescence may differ in these groups could reveal great insights into the function of anuran fluorescent traits. Again, we highlight the importance of testing additional species for the trait of biofluorescence, under a large range of excitation wavelengths, to increase the breadth, depth, and knowledge of this novel trait.

We found evidence for anuran biofluorescence across a broad phylogenetic spectrum and support for all four of Marshall and Johnsen’s (2017) criteria for establishing that a biofluorescent pattern is ecologically significant. Our study supports the idea that anurans are utilizing fluorescent signals as a communication mechanism. The biofluorescence in many frog species matches the perception peak of anuran green rods but strongly differs from background colors reflected during normal frog breeding hours, making biofluorescence most visible during this time. In sum, our results suggest that sensory drive may underlie the evolution of biofluorescence, motivating future research on its function in anuran communication.

## Materials and Methods

### Experimental Design

This study was designed to discover, document, and assess variation in amphibian biofluorescence. To discover fluorescence across the diversity of amphibians, we conducted field surveys of this trait across eight sites representing much of the amphibian biodiversity of lowland South America, the region with the highest species richness of amphibians in the world. While most specimen acquisition was opportunistic via nightly trail surveys, we focused our efforts on collection of anurans, especially treefrogs from the Hylidae family and the *Dendropsophus* genus within this family. We made the choice to focus our sampling on *Dendropsophus* and Hylidae to increase our sample size for future genus- and family-specific studies, balancing the breadth and depth of our data collection. To document biofluorescence, we collected spectrometer recordings and photographs of individual amphibian fluorescence under five excitation sources. To assess variation in the amphibian biofluorescent traits we discovered, we used analysis of variance tests. Specifically, we assessed if biofluorescent emission differed by excitation wavelength to search for evidence of ecological tuning of this novel trait.

### Study sites

We collected data at eight sites spanning four countries across South America. We chose study sites to maximize the diversity of hylid frogs, as preliminary work has shown that this group has the highest presence and diversity of biofluorescence found to date (13–18). We focused efforts on collecting individuals from the genus *Dendropsophus*; our collection sites include the geographic ranges of 77 of the 108 recognized *Dendropsophus* species. These sites include field stations in four South American countries: Colombia, Ecuador, Peru, and Brazil, including four states within Brazil (SP, ES, BA, AM). Data collection occurred from March to May of 2022, within the breeding season of most anurans at these sites. Our field collection schedule was as follows: March 2^nd^-10^th^ at Yasuní Scientific Station, Ecuador; March 13^th^-21^st^ at El Amargal Nature Reserve, Nuquí, Chocó, Colombia; March 24^th^-April 2^nd^ at Los Amigos Conservation Hub (CIRCA), Peru; April 8^th^-13^th^ sampling in Fazenda Michelin, Ubatuba, São Paulo (SP); April 17^th^-20^th^ sampling in Estação Biológica Augusto Ruschi, Aracruz, Espirito Santo (ES); April 23^rd^-27^th^ sampling in Reserva Michelin, Igrapiúna, Bahia (BA); May 2^nd^-5^th^ sampling in Manaus, along Rio Negro via boat, State of Amazonas (AM); May 6^th^-13^th^ sampling in Presidente Figueiredo, AM.

### Field capture and specimen preparation

We collected individuals during night surveys on the trails surrounding each research station we visited. We captured individual amphibians by hand and placed them in a labelled Ziploc plastic bag with air, substrate, and water for transport back to the field station. Each individual was given a field number, and GPS coordinates were taken at the point of capture using a handheld Garmin GPS system. At the field station, all biofluorescent measurements were taken on the same night of capture (see methodology below). The following morning, we euthanized each individual, determined species and sex via dissection, collected tissues, and prepared samples for museum accession. Individual species IDs were determined via region-specific field guides and knowledge of local collaborators. All specimen capture and sample acquisition followed appropriate permit requirements for the specific site (see Acknowledgements).

### Collecting biofluorescence measurements

At the field station, and on the same night of capture, we tested each individual for fluorescence under five different excitation sources spanning a nearly 200nm wavelength range (365-460nm) using the Xite Fluorescent Flashlight System (*NightSea*). The five excitation wavelength ranges were as follows: UV – Ultraviolet (360 – 380nm), VI – Violet (400 – 415nm), RB – Royal blue (440 – 460nm), CY – Cyan (490 – 515nm), and GR – Green (510 – 540nm). We focused on this range of wavelengths to encompass and expand upon wavelengths of biofluorescent excitation used in previous studies (13–18) and to ensure that we did not miss any novel biofluorescent traits. The individual was held by hand beneath the light source, which was suspended above the organism via a *YSLIWC Gooseneck Tripod* stand. We obtained biofluorescent emission spectra for each individual by utilizing a Maya2000 Pro Series UV-VIS Portable Spectrometer with its attached fiber optic cable held above the surface of the amphibian’s skin while beneath the excitation light. This instrument provided information on the emission spectra and intensity of any biofluorescent signals from the frog (200nm-1000nm) in response to each of the five excitation wavelengths. *NightSea* barrier filter glasses matching the respective excitation source were utilized to view and identify potential locations of biofluorescence to test on the body of the amphibian. We utilized the live spectrometer recording acquisition feature of the *OceanView* computer application to determine if a specific location had a fluorescent signal (as defined by the presence of a visible peak at a longer wavelength than the excitation source). If any intensity peak was visible, or if it was questionable if a peak was visible, a spectrometer recording was taken with the probe held approximately two millimeters above that area of skin. We used a 1,000 ms integration time for each spectrometer recording and took three recordings at each location to have technical replicate measurements of the biofluorescent signal at that body location.

In addition, we photographed patterns of biofluorescence under each light source, as well as under a full-spectrum headlamp light source, using a Nikon digital camera and Tiffen camera filters matching the respective excitation source (as specified by *NightSea*). Along with these quantitative measurements, we recorded qualitative descriptions of the color of skin and the color of fluorescence.

### Determining characteristics of biofluorescent measurements

We determined the peak excitation and emission wavelength and intensity from each spectrometer recording and utilized these intensities to calculate a maximum percent biofluorescence emission:

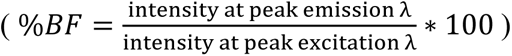

This provided our focal measure of proportion of excitation light re-emitted as fluorescence. All characterizations of spectrometer recordings were completed in R version 4.2.3 (R Core Team 2023). An ASCII file, containing intensity in photon counts for wavelengths ranging 200-1000nm, of each spectrometer recording was saved via the *OceanView* application at time of acquisition and later loaded into R for analysis. Spectra were smoothed via a fifteen-point moving average filter to reduce noise in the spectrum. Utilizing the photobiology package (Aphalo 2015), smoothed spectra were changed to an object of class spectra and normalized to the highest intensity (corresponding to the excitation light) using the normalize () function. We found peak excitation wavelengths by using the which.max() function and confining the parameters to only look for the maximum intensity within the wavelength range of the excitation light source. For example, finding the wavelength that corresponds to the highest intensity (photon count) within the wavelength range of 360-380 nm for UV excitation light, 400-415 nm for VI light, etc. Some of our excitation peak recordings were saturated (maximum >60,000 photon counts). If this was the case, the maximum intensity value available from the spectrometer recording (at 60,000 photons) was utilized. We expect data affected by this saturation would overestimate the intensity of the fluorescence emission recording, though this should not affect the overall significance of our findings because we estimate that saturation occurred at comparable levels across all excitation light sources. All peak emission wavelengths were found using the get_peaks() function from the photobiology package. As a fluorescent signal can produce multiple peaks, the get_peaks() function found all peaks at a wavelength greater than the longest wavelength of the excitation source (greater than 380 nm for UV, 400 nm for VI, etc.). Because fluorescence absorbs light and re-emits it at a longer wavelength, these peaks are all potential fluorescent signals under the respective excitation light. Within the get_peaks() function, we set the ignore_threshold to 0.01 to only collect peaks with a relative size greater than 1% as compared to the tallest (excitation) peak, and we set the span to 100 to define a peak as a datapoint within the wavelength sequence that had an intensity greater than those within a 50-count window on either side of the point. These parameters were chosen via trials adjusting parameters to find peaks of a subset of spectrometer recordings representing the diversity of emission spectra shapes (one emission peak close to the excitation peak, one peak far from excitation, two peaks, no peaks, etc.) and were chosen to reduce the probability that noise in the recording was defined as a peak. Additionally, to further assure that the chosen peaks were significant signals and not noise, any suspected emission peak with an intensity greater than the excitation peak was removed, as this was likely a sign of an inaccurate spectrometer recording. In these cases, and in any cases where no peaks were found, “NA” was input for the peak wavelength and intensity values. The intensity at each peak emission wavelength was divided by the intensity at the peak excitation wavelength for the respective spectrometer recording to calculate the average percent of the excitation light that is re-emitted as fluorescence.

As stated, we took three spectrometer recordings at each location on each individual to have technical replicate measurements of the biofluorescent signal at that body location. The calculated percentages of biofluorescence emission for these three recordings were averaged for each body location. Because our data set contained spectrometer recordings from a different number of body locations, and often different specific body regions, from each individual (due to the variation in physical biofluorescent patterns that we were attempting to capture across a wide range of species), we only used the maximum percent biofluorescence emission (from any body region, under *each* of the five excitation light sources) for downstream analyses. Hence, for all analyses of variance, the unit of measurement used was the maximum percent of biofluorescence emission recorded from each individual.

We also examined the body location from which the maximum biofluorescent recording from each individual was taken. For this, we used the maximum percent biofluorescence emission (from any body region, under *any* light source). Utilizing the maximum fluorescent emission recording for each individual insured that only one body location per individual was being considered, although, examining the maximum biofluorescent emission recording for each individual under each excitation light source (as in the analyses above) produced similar percentages of fluorescence by body location. For easier comparison, the body regions were summarized into the following nine groups: cloaca, dorsal (including spectrometer recordings with a body location specified from any dorsal pattern), eye, facial pattern (including lip, spots under the eye, snout, etc.), flank, inguinal region, limb (including forelimbs, thigh, etc.), throat (including vocal sac), and ventral (including any ventral pattern). The percentage of maximum biofluorescent recordings from each of the nine body locations were calculated for each light source.

### Statistical Analysis

i. *Evaluate Criterion 1:* The fluorescent pigment will absorb the dominant wavelengths of the environment.

To evaluate evidence for Criterion 1, we determined which excitation source produced the maximum fluorescent signal and evaluated the distance of that excitation wavelength to the dominant wavelength of the twilight environment. The excitation wavelength that produces fluorescence is the wavelength of light absorbed by the fluorescent pigment. The residuals of the individual maximum percent of biofluorescence emission recordings were non-normal, hence, we utilized the non-parametric Kruskal-Wallis test in place of a one-way analysis of variance to determine if the percent of biofluorescent emission differed by excitation light source. Additionally, we utilized Pairwise Dunn’s tests with Holm adjustment to determine which excitation light differed from each other in biofluorescent emission.

Similarly, we utilized a non-parametric Kruskal-Wallis test and Pairwise Dunn’s tests with Holm adjustment to determine if the distance to the dominant wavelength of the twilight environment differed by excitation light source. We calculated this distance by subtracting 457.5 nm (the median dominant wavelength in the twilight environment (27)) from the peak excitation wavelength value obtained for each individual (as described above).

ii. *Evaluate Criterion 2:* The fluorescence will be viewed against a contrasting background.

To evaluate evidence for Criterion 2, we assessed if the peak emission wavelength of the blue-light-induced fluorescence matched the least dominant wavelengths of the twilight environment better than expected by chance. Fluorescent emission wavelengths that match the least dominant wavelengths of light at twilight will produce the most contrast with the background environment.

As above, we utilized a non-parametric Kruskal-Wallis one-way analysis of variance test and Pairwise Dunn’s tests with Holm adjustment to determine if the wavelength of biofluorescent emission differed by excitation light source. Within the biofluorescent emission produced by the blue excitation light source there were two dominant emission peaks: a “green” peak and an “orange” peak (Fig. 4). We utilized a randomization test to assess if each of these emission peaks matched the least dominant wavelengths of the twilight environment better than expected by chance. A “green” peak was defined as a wavelength less than 550nm and an “orange” peak defined as a wavelength greater than 550nm. For the randomization test, we determined the average value of twilight irradiance at the emission wavelengths of that peak (“green” or “orange”). This is the test statistic whose value was 56.38 and 38.72 relative twilight irradiance for the green and orange peaks respectively. We then took a sample of the same size (192 and 316 individuals for “green” and “orange” peaks respectively) and randomly selected a wavelength between 400 and 700 nm, then calculated a new value for the test statistic for those wavelengths. We repeated this for 10,000 iterations and compared the test statistics in the randomization distribution to the observed test statistic value to determine if the peak fluorescent emission matched the wavelengths least available in the twilight environment better than by chance (α = 0.05).

iii. *Evaluate Criterion 3:* Organisms viewing the fluorescence will have spectral sensitivity in the fluorescent emission range.

To evaluate evidence for Criterion 3, we assessed if the peak emission wavelength of the blue-light-induced fluorescence matched the peak sensitivity of the green sensitive anuran rod better than expected by chance. As above, we utilized a randomization test to assess if each of the “green” and “orange” emission peaks matched the peak sensitivity of the green sensitive anuran rod better than expected by chance. We determined the average optical sensitivity at the emission wavelengths of that peak (“green” or “orange”). This is the test statistic whose value was 92.84 and 3.18 relative rod sensitivity for the green and orange peaks respectively. We then took a sample of the same size (192 and 316 individuals for “green” and “orange” peaks respectively) and randomly selected a wavelength between 400 and 700 nm, then calculated the value of the test statistic for those wavelengths. We repeated this for 10,000 iterations and compared the null distribution to the observed test statistic value to determine if the peak fluorescent emission matched the wavelengths of greatest sensitivity in the green sensitive rod better than by chance (α = 0.05).

Considering that both Criterion 2 and Criterion 3 need to be met for ecological significance, we compared the blue-light-induced “green” emission peak to the tradeoff spectrum of twilight irradiance and anuran rod sensitivity, as follows. We divided the green-sensitive rod spectrum (28) by the twilight irradiance spectrum (27) to obtain the tradeoff spectrum. Both the twilight and rod spectra were standardized before this calculation. As above, we then utilized a randomization test to assess if the “green” emission peak matched the tradeoff spectrum better than expected by chance. We calculated the average tradeoff value at the emission wavelengths of the “green” peak. This is the test statistic. We then took a sample of the same size (192 individuals) and randomly selected a wavelength between 400 and 700 nm, then calculated the value of the test statistic for those wavelengths. We repeated this for 10,000 iterations and compared the test statistic values in the randomization distribution to the observed test statistic to determine if the “green” peak fluorescent emission maximized the wavelengths to meet both Criterion 2 and Criterion 3 better than by chance.

For all randomization tests, we ran a sensitivity analysis to determine at which effective sample size our statistical interpretations held. To do so, we changed the sample size of the randomization distribution and determined at which sample size, n, the null distribution of test statistic values was not significantly different than the observed test statistic, as defined above at α = 0.05. The effective sample size for all significant randomization tests was n = 2, except for that test comparing the “green” peak fluorescent emission and twilight irradiance spectrum whose effective sample size was n = 13. Hence, our statistical interpretations stated in the Results section above hold as long as our data contains at least two and thirteen independent groups respectively. As we had representatives from 82 species within our sample size of 192 “green” peak individuals, this effective sample size is very likely, despite the unknown influence of phylogeny on anuran biofluorescence.

## Acknowledgments

We would like to thank everyone who made this work possible, including all field stations at which data collection took place: Yasuni Scientific Station, El Amargal Nature Reserve, Los Amigos Conservation Hub (CIRCA), Fazenda Michelin, Estação Biológica Augusto Ruschi, Reserva Michelin, and Tartarugas da Amazonia (and surrounding indigenous communities including the Nova Esperança community of the Baré ethnic group and the Saracá community). Additionally, we thank Rafael Benettii, Lucas Neves, Victor Araújo, Thalia Corahua, and Schyler Ellsworth for their help in the field and with sample preparation. Finally, we would like to thank Carlos Taboada for his advice on appropriate equipment and methodology during field expedition planning.

Collections in Ecuador were made under the authority of the *Ministerio de Ambiente, Agua y Transición Ecológica* No. 2034. Vouchers were deposited in The Zoology Museum at the Pontifical Catholic University of Ecuador (*QCAZ*). Collections in Peru were made under the authority of the *Ministerio de Agricultura y Riego* No. 2022-0000809. Vouchers were deposited in the *Museo de Historia Natural San Marcos* (*UNMSM*). Collections in Colombia were made under the authority of the *Permiso marco* No. 1177 of 09 October 2014 granted by the *Autoridad Nacional de Licencias Ambientales* (ANLA) to the Universidad de los Andes (UniAndes). Relevant animal care and use protocols for amphibians were approved by the CICUA of UniAndes under POE 22-001 and vouchers were deposited in the Museo de Historia Natural C.J. Marinkelle at UniAndes. Collections in Brazil were made under the authority of the *Brazil Instituto Chico Mendes de Conservação da Biodiversidade* (*ICMBio*) via collaboration with the Instituto Butantan. Vouchers were deposited at the Instituto Butantan and registered in the *SisGen* system by time of publication. All data collection methods were approved by the IACUC under protocol IPROTO202100000007.

## Funding

The Lamarr and Edith Trott Scholarship (CW)

The Explorers Club Rolex Grant (CW)

Horace Loftin Endowment (CW)

Martin Family Graduate Fellowship in Biological Sciences (CW)

Society for Systematic Biologists Graduate Student Research Award (CW)

Robert B. Short Scholarship in Zoology (CW)

## Author contributions

Conceptualization: CW, SRR, FAV, RMB

Methodology: CW, SRR, FAV, AC, VHA, EF, FG, RMB, AL

Investigation: CW, SRR, FAV, RMB Visualization: CW, AL

Supervision: EML Writing—original draft: CW

Writing—review & editing: CW, SRR, FAV, AC, VHA, EF, FG, RMB, AL, EML

## Data and materials availability

All code will be deposited to Dryad (https://datadryad.org) at time of submission. All specimen data can be accessed via the museum/institution in which the individual was accessioned (see Acknowledgement section above). All spectrometer recordings and photographs are available by reasonable request. All other data are available in the main text or the supplementary materials.

