## Supplementary Materials for "Evidence for ecological tuning of novel anuran biofluorescent signals"

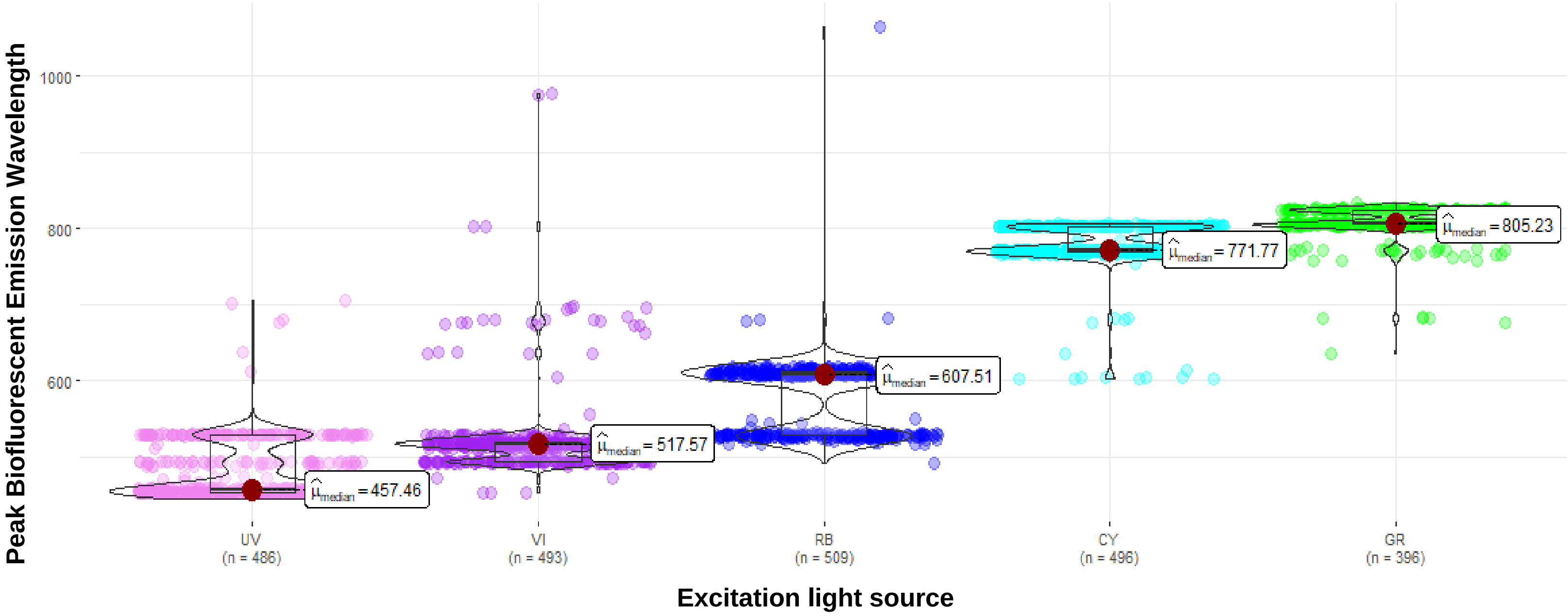


**Fig. S1. Peak biofluorescent emission by excitation source.** The wavelength of biofluorescent emission by excitation light source: UV – Ultraviolet (360-380 nm), VI – Violet (400-415 nm), RB – Royal blue (440-460 nm), CY – Cyan (490-515 nm), and GR – Green (510-540 nm). Each point represents one individual (the wavelength of the maximum percent biofluorescent emission recorded for that individual under that excitation light source).

**Table S1. Maximum percent biofluorescent emission by family.** The maximum percent of biofluorescence emission measurements from each individual under each light source collected in this study presented by family. The excitation light sources are as follows: UV – Ultraviolet (360-380 nm), VI – Violet (400-415 nm), RB – Royal blue (440-460 nm), CY – Cyan (490-515 nm), and GR – Green (510-540 nm). These data are plotted in Fig. 1.

| Family | Excitation | n | min | max | median | iqr | mean | sd | se | ci |
|---|---|---|---|---|---|---|---|---|---|---|
| *Aromobatidae* | UV | 3 | 4.416 | 12.981 | 5.687 | 4.283 | 7.695 | 4.622 | 2.669 | 11.482 |
| *Brachycephalidae* | UV | 1 | 4.598 | 4.598 | 4.598 | 0 | 4.598 | NA | NA | NA |
| *Bufonidae* | UV | 19 | 3.773 | 13.343 | 4.984 | 1.518 | 5.835 | 2.555 | 0.586 | 1.231 |
| *Caeciliidae* | UV | 1 | 9.661 | 9.661 | 9.661 | 0 | 9.661 | NA | NA | NA |
| *Centrolenidae* | UV | 7 | 3.431 | 32.436 | 6.876 | 4.982 | 10.839 | 9.92 | 3.749 | 9.175 |
| *Craugastoridae* | UV | 24 | 3.088 | 14.454 | 5.549 | 5.14 | 6.481 | 3.034 | 0.619 | 1.281 |
| *Dendrobatidae* | UV | 4 | 4.497 | 8.325 | 4.958 | 1.385 | 5.685 | 1.789 | 0.895 | 2.847 |
| *Eleutherodactylidae* | UV | 6 | 3.633 | 7.928 | 4.6 | 2.746 | 5.263 | 1.829 | 0.747 | 1.919 |
| *Hylidae* | UV | 324 | 2.345 | 77.223 | 5.667 | 3.744 | 7.744 | 7.184 | 0.399 | 0.785 |
| *Leptodactylidae* | UV | 36 | 3.285 | 19.767 | 6.306 | 3.231 | 6.918 | 3.421 | 0.57 | 1.158 |
| *Microhylidae* | UV | 13 | 3.6 | 21.356 | 7.638 | 7.057 | 8.997 | 5.267 | 1.461 | 3.183 |
| *Odontophrynidae* | UV | 4 | 4.68 | 10.031 | 4.777 | 1.354 | 6.066 | 2.643 | 1.322 | 4.206 |
| *Pipidae* | UV | 1 | 5.41 | 5.41 | 5.41 | 0 | 5.41 | NA | NA | NA |
| *Strabomantidae* | UV | 43 | 2.968 | 31.69 | 5.639 | 4.255 | 8.005 | 5.46 | 0.833 | 1.68 |
| *Aromobatidae* | VI | 3 | 4.133 | 8.147 | 4.634 | 2.007 | 5.638 | 2.187 | 1.263 | 5.433 |
| *Brachycephalidae* | VI | 1 | 8.108 | 8.108 | 8.108 | 0 | 8.108 | NA | NA | NA |
| *Bufonidae* | VI | 21 | 3.901 | 40.494 | 6.636 | 7.442 | 12.268 | 10.283 | 2.244 | 4.681 |
| *Caeciliidae* | VI | 1 | 6.77 | 6.77 | 6.77 | 0 | 6.77 | NA | NA | NA |
| *Centrolenidae* | VI | 7 | 5.327 | 20.754 | 10.744 | 4.983 | 10.784 | 5.167 | 1.953 | 4.779 |
| *Craugastoridae* | VI | 24 | 3.857 | 40.719 | 16.958 | 10.48 | 17.96 | 9.607 | 1.961 | 4.057 |
| *Dendrobatidae* | VI | 4 | 9.67 | 15.055 | 12.028 | 3.834 | 12.195 | 2.589 | 1.295 | 4.12 |
| *Eleutherodactylidae* | VI | 6 | 10.331 | 34.732 | 13.421 | 9.089 | 17.46 | 9.56 | 3.903 | 10.032 |
| *Hylidae* | VI | 325 | 2.063 | 94.916 | 7.861 | 9.971 | 14.454 | 15.584 | 0.864 | 1.701 |
| *Leptodactylidae* | VI | 35 | 3.299 | 79.262 | 9.805 | 11.799 | 15.057 | 14.196 | 2.4 | 4.876 |
| *Microhylidae* | VI | 15 | 7.884 | 49.271 | 12.081 | 10.951 | 17.908 | 12.439 | 3.212 | 6.889 |
| *Odontophrynidae* | VI | 4 | 5.375 | 9.345 | 7.521 | 3.553 | 7.441 | 2.14 | 1.07 | 3.405 |
| *Pipidae* | VI | 1 | 18.682 | 18.682 | 18.682 | 0 | 18.682 | NA | NA | NA |
| *Strabomantidae* | VI | 46 | 3.624 | 36.335 | 12.805 | 13.547 | 15.324 | 8.897 | 1.312 | 2.642 |
| *Aromobatidae* | RB | 3 | 3.932 | 39.744 | 5.653 | 17.906 | 16.443 | 20.197 | 11.661 | 50.173 |
| *Brachycephalidae* | RB | 1 | 7.121 | 7.121 | 7.121 | 0 | 7.121 | NA | NA | NA |
| *Bufonidae* | RB | 24 | 4.117 | 16.213 | 7.113 | 5.519 | 8.357 | 3.636 | 0.742 | 1.535 |
| *Caeciliidae* | RB | 1 | 20.554 | 20.554 | 20.554 | 0 | 20.554 | NA | NA | NA |
| *Centrolenidae* | RB | 7 | 9.427 | 35.346 | 14.069 | 5.036 | 17.284 | 8.538 | 3.227 | 7.897 |
| *Craugastoridae* | RB | 26 | 4.521 | 84.697 | 16.12 | 27.148 | 27.498 | 23.219 | 4.554 | 9.378 |
| *Dendrobatidae* | RB | 5 | 4.043 | 30.045 | 9.109 | 4.223 | 13.103 | 10.019 | 4.481 | 12.441 |
| *Eleutherodactylidae* | RB | 6 | 8.789 | 27.566 | 18.748 | 13.244 | 17.748 | 8.164 | 3.333 | 8.567 |
| *Hylidae* | RB | 331 | 2.646 | 96.85 | 10.706 | 11.288 | 15.992 | 15.378 | 0.845 | 1.663 |
| *Leptodactylidae* | RB | 37 | 3.754 | 60.841 | 16.057 | 14.299 | 19.328 | 13.549 | 2.228 | 4.518 |
| *Microhylidae* | RB | 15 | 9.521 | 40.801 | 27.759 | 18.433 | 26.954 | 10.397 | 2.685 | 5.758 |
| *Odontophrynidae* | RB | 4 | 5.033 | 12.466 | 7.152 | 1.945 | 7.951 | 3.172 | 1.586 | 5.047 |

| | | | | | | | | | | |
|---|---|---|---|---|---|---|---|---|---|---|
| *Pipidae* | RB | 1 | 9.141 | 9.141 | 9.141 | 0 | 9.141 | NA | NA | NA |
| *Plethodontidae* | RB | 2 | 10.485 | 19.059 | 14.772 | 4.287 | 14.772 | 6.063 | 4.287 | 54.471 |
| *Strabomantidae* | RB | 46 | 4.876 | 60.381 | 16.063 | 21.1 | 20.704 | 14.27 | 2.104 | 4.238 |
| *Aromobatidae* | CY | 3 | 3.661 | 5.046 | 3.939 | 0.693 | 4.215 | 0.733 | 0.423 | 1.821 |
| *Brachycephalidae* | CY | 1 | 6.385 | 6.385 | 6.385 | 0 | 6.385 | NA | NA | NA |
| *Bufonidae* | CY | 23 | 2.919 | 80.253 | 5.176 | 2.639 | 8.923 | 15.689 | 3.271 | 6.784 |
| *Caeciliidae* | CY | 1 | 8.507 | 8.507 | 8.507 | 0 | 8.507 | NA | NA | NA |
| *Centrolenidae* | CY | 7 | 3.664 | 16.618 | 6.893 | 3.213 | 7.364 | 4.407 | 1.666 | 4.076 |
| *Craugastoridae* | CY | 26 | 3.425 | 10.696 | 6.61 | 3.129 | 6.793 | 2.013 | 0.395 | 0.813 |
| *Dendrobatidae* | CY | 3 | 4.252 | 9.534 | 7.256 | 2.641 | 7.014 | 2.649 | 1.53 | 6.581 |
| *Eleutherodactylidae* | CY | 6 | 4.776 | 14.273 | 7.314 | 2.909 | 8.165 | 3.411 | 1.392 | 3.58 |
| *Hylidae* | CY | 327 | 2.175 | 22.994 | 5.746 | 4.074 | 7.114 | 3.899 | 0.216 | 0.424 |
| *Leptodactylidae* | CY | 33 | 3.154 | 23.897 | 6.51 | 4.943 | 8.043 | 4.919 | 0.856 | 1.744 |
| *Microhylidae* | CY | 15 | 4.018 | 19.42 | 9.386 | 3.198 | 9.125 | 3.661 | 0.945 | 2.027 |
| *Odontophrynidae* | CY | 4 | 4.446 | 19.675 | 6.244 | 4.374 | 9.152 | 7.073 | 3.537 | 11.255 |
| *Pipidae* | CY | 1 | 3.339 | 3.339 | 3.339 | 0 | 3.339 | NA | NA | NA |
| *Plethodontidae* | CY | 1 | 8.659 | 8.659 | 8.659 | 0 | 8.659 | NA | NA | NA |
| *Strabomantidae* | CY | 45 | 3.824 | 32.45 | 6.599 | 3.596 | 7.918 | 4.877 | 0.727 | 1.465 |
| *Aromobatidae* | GR | 3 | 4.412 | 7.34 | 5.925 | 1.464 | 5.892 | 1.464 | 0.845 | 3.638 |
| *Brachycephalidae* | GR | 1 | 6.836 | 6.836 | 6.836 | 0 | 6.836 | NA | NA | NA |
| *Bufonidae* | GR | 15 | 3.152 | 12.934 | 4.078 | 0.873 | 5.08 | 2.604 | 0.672 | 1.442 |
| *Caeciliidae* | GR | 1 | 9.925 | 9.925 | 9.925 | 0 | 9.925 | NA | NA | NA |
| *Centrolenidae* | GR | 6 | 6.792 | 13.748 | 10.658 | 3.024 | 10.241 | 2.542 | 1.038 | 2.667 |
| *Craugastoridae* | GR | 22 | 3.813 | 18.779 | 7.134 | 3.932 | 7.825 | 3.832 | 0.817 | 1.699 |
| *Dendrobatidae* | GR | 2 | 7.96 | 40.978 | 24.469 | 16.509 | 24.469 | 23.347 | 16.509 | 209.765 |
| *Eleutherodactylidae* | GR | 2 | 3.861 | 4.186 | 4.023 | 0.163 | 4.023 | 0.23 | 0.163 | 2.069 |
| *Hylidae* | GR | 277 | 2.756 | 64.78 | 6.947 | 4.082 | 8.216 | 5.291 | 0.318 | 0.626 |
| *Leptodactylidae* | GR | 25 | 3.751 | 29.034 | 6.808 | 4.047 | 8.501 | 5.872 | 1.174 | 2.424 |
| *Microhylidae* | GR | 10 | 1.948 | 16.697 | 5.047 | 2.464 | 6.796 | 4.531 | 1.433 | 3.241 |
| *Odontophrynidae* | GR | 4 | 3.606 | 6.169 | 5.559 | 1.285 | 5.223 | 1.17 | 0.585 | 1.861 |
| *Pipidae* | GR | 1 | 7.833 | 7.833 | 7.833 | 0 | 7.833 | NA | NA | NA |
| *Strabomantidae* | GR | 27 | 2.256 | 13.467 | 6.661 | 2.637 | 6.841 | 2.327 | 0.448 | 0.92 |

**Table S2. Pair-wise comparison of distance to dominant wavelength of twilight by excitation light source.** Dunn (1964) multiple comparison p-values adjusted with the Holm method for each excitation light source wavelength: UV – Ultraviolet (360-380 nm), VI – Violet (400-415 nm), RB – Royal blue (440-460 nm), CY – Cyan (490-515 nm), and GR – Green (510-540 nm). (*) Indicates significance at alpha = 0.05. All pairwise comparisons of distance to dominant twilight wavelength are significantly different. blue (440-460 nm) excitation light was closest to the dominant twilight wavelength of 457.5 nm (27).

| | **UV** | **CY** | **GR** | **VI** |
|---|---|---|---|---|
| **CY** | -32.000<br>4.330e-224* | | | |
| **GR** | -20.219<br>1.661e-90* | -9.997<br>1.577e-23* | | |
| **VI** | 11.508<br>1.808e-30* | -20.549<br>2.360e-93* | -9.383<br>3.205e-21* | |
| **RB** | -43.934<br>0.000e+00* | 11.790<br>8.784e-32* | 21.155<br>8.639e-99* | -32.453<br>2.216e-230* |

**Table S3. Maximum biofluorescent emission recording count by body location and excitation light source.** The body location from which the maximum biofluorescent emission recording from each individual was taken, separated by excitation light source (the maximum fluorescent emission recording under *any* light source). The excitation light sources are as follows: UV – Ultraviolet (360-380 nm), VI – Violet (400-415 nm), RB – Royal blue (440-460 nm), CY – Cyan (490-515 nm), and GR – Green (510-540 nm). Body regions were summarized into the following nine groups: cloaca, dorsal (including spectrometer recordings with a body location specified from any dorsal pattern), eye, facial pattern (including lip, spots under the eye, snout, etc.), flank, inguinal region, limb (including forelimb, thigh, etc.), throat (including vocal sac), and ventral (including any ventral pattern). The percentage of maximum biofluorescent emission recordings from each of the nine body locations were calculated for each light source.

| **Body Location** | **Count** | **Percent (%)** | **Excitation** |
|---|---|---|---|
| dorsal | 8 | 26 | UV |
| eye | 2 | 6 | UV |
| flank | 1 | 3 | UV |
| inguinal region | 1 | 3 | UV |
| throat | 12 | 39 | UV |
| ventral | 7 | 23 | UV |
| cloaca | 1 | 1 | VI |
| dorsal | 42 | 27 | VI |
| eye | 1 | 1 | VI |
| facial pattern | 5 | 3 | VI |
| flank | 23 | 15 | VI |
| inguinal region | 4 | 3 | VI |
| limb | 7 | 4 | VI |
| throat | 40 | 26 | VI |
| ventral | 33 | 21 | VI |
| dorsal | 46 | 18 | RB |
| eye | 2 | 1 | RB |
| facial pattern | 4 | 2 | RB |
| flank | 30 | 11 | RB |
| inguinal region | 13 | 5 | RB |
| limb | 8 | 3 | RB |
| throat | 74 | 28 | RB |
| ventral | 84 | 32 | RB |
| dorsal | 9 | 32 | CY |
| eye | 1 | 4 | CY |
| flank | 2 | 7 | CY |
| throat | 5 | 18 | CY |
| ventral | 11 | 39 | CY |
| dorsal | 11 | 31 | GR |
| flank | 4 | 11 | GR |
| inguinal region | 5 | 14 | GR |
| throat | 11 | 31 | GR |
| ventral | 5 | 14 | GR |

**Table S4. Maximum biofluorescent emission recording under each excitation light source by body location.** The body location from which the maximum biofluorescent emission recording from each individual, under each excitation light source, was taken (the maximum fluorescent emission recording under *each* light source). The excitation light sources are as follows: UV – Ultraviolet (360-380 nm), VI – Violet (400-415 nm), RB – Royal blue (440-460 nm), CY – Cyan (490-515 nm), and GR – Green (510-540 nm). Body regions were summarized into the following nine groups: cloaca, dorsal (including spectrometer recordings with a body location specified from any dorsal pattern), eye, facial pattern (including lip, spots under the eye, snout, etc.), flank, inguinal region, limb (including forelimb, thigh, etc.), throat (including vocal sac), and ventral (including any ventral pattern).

| Body Location | Count | Percent (%) | Excitation |
|---|---|---|---|
| cloaca | 2 | <1 | UV |
| dorsal | 76 | 16 | UV |
| eye | 4 | 1 | UV |
| facial pattern | 2 | <1 | UV |
| flank | 38 | 8 | UV |
| inguinal region | 12 | 2 | UV |
| limb | 27 | 6 | UV |
| throat | 231 | 48 | UV |
| ventral | 94 | 19 | UV |
| cloaca | 6 | 1 | VI |
| dorsal | 162 | 33 | VI |
| eye | 4 | 1 | VI |
| facial pattern | 7 | 1 | VI |
| flank | 68 | 14 | VI |
| inguinal region | 18 | 4 | VI |
| limb | 31 | 6 | VI |
| throat | 117 | 24 | VI |
| ventral | 79 | 16 | VI |
| dorsal | 112 | 22 | RB |
| eye | 4 | 1 | RB |
| facial pattern | 6 | 1 | RB |
| flank | 58 | 11 | RB |
| inguinal region | 18 | 4 | RB |
| limb | 18 | 4 | RB |
| throat | 150 | 30 | RB |
| ventral | 142 | 28 | RB |
| cloaca | 1 | <1 | CY |
| dorsal | 112 | 23 | CY |
| eye | 1 | <1 | CY |
| facial pattern | 5 | 1 | CY |
| flank | 35 | 7 | CY |
| inguinal region | 15 | 3 | CY |
| limb | 14 | 3 | CY |
| throat | 180 | 36 | CY |
| ventral | 132 | 27 | CY |
| cloaca | 1 | <1 | GR |
| dorsal | 100 | 25 | GR |
| facial pattern | 4 | 1 | GR |
| flank | 36 | 9 | GR |
| inguinal region | 21 | 5 | GR |
| limb | 6 | 2 | GR |
| throat | 140 | 35 | GR |
| ventral | 88 | 22 | GR |
